# Activated NK cells with a predominance of inhibitory receptors and a decidual-like phenotype expand after autologous HSCT in children with tumors

**DOI:** 10.1101/2024.07.01.601507

**Authors:** Gabirel Astarloa-Pando, Diego Polanco-Alonso, Víctor Sandá, Ane Orrantia, Ainhoa Amarilla-Irusta, Silvia Pérez-Fernández, Raquel Pérez-Garay, Arrate Sevilla, Ainara Lopez-Pardo, Aritz Tijero, José J. Úriz, Mercedes Rey, Itziar Astigarraga, Bárbara Manzanares-Martin, Naiara G. Bediaga, Laura Amo, Olatz Zenarruzabeitia, Francisco Borrego

**Author notes:** Correspondence: Francisco Borrego MD, PhD. Immunopathology Group, Biobizkaia Health Research Institute, 48903 Barakaldo, Spain.

## Abstract

Early immune reconstitution after autologous hematopoietic stem cell transplantation (autoHSCT) is associated with a better outcome in a variety of cancers. NK cells constitute the first lymphocyte subset to recover in the blood after autoHSCT. We have in-depth characterized them in pediatric patients with different tumors and found that, immediately after autoHSCT, NK cells transiently acquired a decidual-like phenotype, were more immature and activated, and exhibited an increased expression of inhibitory receptors, while activating receptors levels were diminished. This activated and decidual-like phenotype was characterized by increased CD56, CD9, CD49a, CD151, CD38, HLA-DR and CD55 expression. We also determined plasma levels of several cytokines and found that their concentrations were associated with the observed changes in NK cells phenotype. *In vitro* experiments, including flow cytometry and single-cell RNA sequencing (scRNA-seq), recapitulated the changes observed in NK cells early after autoHSCT. Specifically, results revealed that the combination of IL-15 and TGF-β induced, at least partially, this distinctive phenotype on NK cells after autoHSCT. Finally, we have observed a positive correlation between relapse and the percentage of CD56^dim^ NK cells shortly after autoHSCT in our cohort of pediatric patients. Altogether, our work is of relevance to understand the physiopathology of NK cells during the immune system reconstitution after autoHSCT in children and potentially help in the management of these patients.

## INTRODUCTION

Autologous hematopoietic stem cell transplantation (autoHSCT) is a widespread procedure used to treat many different types of cancer, including hematological tumors in adults and solid tumors in children. It is a relatively common therapeutic tool to combat not only hematological malignancies, such as multiple myeloma (MM) or non-Hodgkin lymphoma (NHL), but also for some pediatric solid tumors such as neuroblastoma (NBL), and might be a salvage option for other tumors like lymphomas and sarcomas ^1–3^. Early immune reconstitution following autoHSCT is associated with a better outcome in a variety of cancers ^4–6^. Day 15 absolute lymphocyte count (ALC-15) of ≥ 500 cells/µl after autoHSCT is an independent prognostic indicator in MM and NHL patients ^5^ and natural killer (NK) cells are a relevant lymphocyte subset in ALC-15 that affects the outcome following autoHSCT ^7–9^, as they are the first lymphocyte subset reaching normal circulating levels after HSCT ^10–12^.

NK cells can directly kill target cells through different mechanisms and they have a central role in the defense against viral-infected and cancer cells ^13–16^. Human circulating NK cells have been classified in two major subsets with different functionalities, depending on the expression of CD56 and CD16: CD56^bright^CD16^low/−^ (CD56^bright^) and CD56^dim^CD16+ (CD56^dim^). CD56^bright^ NK cells produce large amounts of immunomodulatory cytokines and chemokines but have low cytotoxicity unless they are activated by cytokines. On the other hand, CD56^dim^ NK cells are more cytotoxic but produce lower amounts of cytokines ^17–20^. NK cells express activating and inhibitory receptors whose integrated signals will determine their response ^21^. In this manner, they do not get activated when they recognize autologous determinants such as human class I human leukocyte antigens (HLA-I) through inhibitory killer immunoglobulin-like receptors (KIRs) and CD94/NKG2A receptors. On the other hand, NK cells are activated when they interact with virus-infected or cancer cells, either by the direct detection of upregulated stress-induced self-molecules, or via antibody-dependent cell-mediated cytotoxicity (ADCC) ^16,19,22,23^. Given the potent anti-tumor effect of NK cells and their early recovery after autoHSCT, studying their biology and reconstitution in this setting is of utmost interest. Studies are scarce, and they are mostly focused on adult patients. They show that following autoHSCT, and shortly after leukocyte recovery, there is a redistribution of NK cells subsets. For example, in patients with MM and lymphoma it has been described that there is an increased frequency of immature CD56^bright^NKG2A+ NK cells, expressing high levels of KIRs and CD57, early after transplant ^24^. Another MM study showed a similar redistribution of NK cell populations characterized by a high proliferative rate and an increase in the frequency of both CD56^bright^ NK cells as well as the most immature population within the CD56^dim^ NK subset (CD57-NKG2A+) ^25^. In general, NK cell function recovers early after autoHSCT ^24,25^ and transcriptomic analyses have revealed profound changes related, among others, to cell cycle, DNA replication and the mevalonate pathway ^26^. Interestingly, following autoHSCT in MM patients there is an expansion of a CD9+ decidual-like NK cell subset that is characterized by high granzyme B and perforin expression levels ^26^. Nevertheless, the factors responsible for this CD9+ NK cell subset expansion are not known.

In addition, it has been described that the number of NK cells and the frequency of several subsets are associated with survival ^12,27^. For example, it has been shown that MM patients with lower frequencies of the mature highly differentiated NKG2A-CD57+ NK cell subset after autoHSCT had a better outcome ^25^. Other researchers have found that MM patients in long-term complete response after autoHSCT exhibit a higher frequency of NK cells expressing the inhibitory receptors KIR2DL1 and NKG2A and reduced numbers of NKp46+ NK cells ^28^. Interestingly, as in allogeneic HSCT (alloHSCT), there could be a NK cell-mediated graft-versus-tumor (GvT) effect in autoHSCT, that is related to the inhibitory KIR - HLA-I receptor-ligand mismatch and affinity interactions between them ^29–33^. Indeed, KIR and HLA-I genotypes have shown to have a significant influence in NBL patients undergoing autoHSCT and anti-GD2 treatment ^30,34^.

Despite the advances in adults, in pediatric patients there is still a substantial need to understand the complexity of the immune system reconstitution after autoHSCT. In this context, an in depth characterization of the NK cell lymphocyte subset, exploring the biology and physiopathology of NK cells in pediatric patients, would be of great value. In the present study, we have found that following autoHSCT there are significant changes in the expression of inhibitory and activating receptors, as well as in activation markers, together with a transient expansion of NK cells with a decidual-like phenotype. Furthermore, plasma levels of soluble molecules that are relevant for NK cell differentiation, survival and function were analyzed. Changes in the levels of interleukin 15 (IL-15) and transforming growth factor beta 1 (TGF-β1 or TGF-β hereafter) may explain the acquisition of the decidual-like phenotype. Also, analyses of maturation markers and KIR expression indicate that NK cell education and maturation are uncoupled processes. Altogether, our findings can be of relevance to understand the biology and physiopathology of NK cells during their reconstitution following autoHSCT and, therefore, help to design better therapies for children with cancer.

## MATERIALS AND METHODS

### Patients’ characteristics and study design

Blood samples from 13 children diagnosed with different cancers and that received autoHSCT were collected for this study. Patients’ clinical features are described in Table 1. Samples were obtained at five different time points: pre-transplantation (S1), after reaching leucocyte recovery: more than 1000 leukocytes/µl, usually around day 12 after autoHSCT (S2), 30 days after autoHSCT (S3), 100 days after autoHSCT (S4) and 180 days after autoHSCT (S5). Blood samples from adult healthy donors were also collected. Sample collection was carried out through the Basque Biobank for Research (https://www.biobancovasco.bioef.eus), which complies with the quality management, traceability and biosecurity, set out in the Spanish Law 14/2007 of Biomedical Research and in the Royal Decree 1716/2011. The study was approved by the Basque Ethics Committee for Clinical Research (BIO14/TP/003, PI+CES+INC-BIOEF 2017-03) and the Cruces University Hospital Ethics Committee (E23/78). All subjects provided written and signed informed consent in accordance with the Declaration of Helsinki.

**Table 1.** Patientś clinical characteristics.

|  |  | **n (%)** |
| --- | --- | --- |
| **Gender** | Male | 8 (61.5%) |
|  | Female | 5 (38.5%) |
| **Cancer type<sup>a</sup>** | Neuroblastoma | 4 (30.8%) |
|  | Other tumors | 9 (69.2%) |
| **Mobilization regimen** | G-CSF | 13 (100%) |
| **Conditioning regimen** | Busulfan-melphalan | 4 (30.8%) |
|  | Melphalan | 3 (23.1%) |
|  | Thiotepa + cyclophosphamide | 2 (15.4%) |
|  | Other | 4 (30.8%) |
| **Second HSCT** | Yes | 5 (38.5%) |
|  | No | 8 (61.5%) |
| **Complementary treatment** | Radiotherapy | 3 (23.1%) |
|  | G-CSF | 2 (15.4%) |
|  | G-CSF ± opioids ± antibiotics | 5 (38.5%) |
|  | No | 3 (23.1%) |
| **Complications after HSCT** | Yes | 12 (92.3%) |
|  | No | 1 (7.7%) |
| **Maintenance regimen** | Anti-GD2 + cis-retinoic acid | 3 (23.1%) |
|  | Anti-GD2 + rapamycin | 1 (7.7%) |
|  | Lenalidomide + bortezomib | 1 (7.7%) |
|  | No | 8 (61.5%) |
| **Relapse** | Yes | 6<sup>b</sup> (46.2%) |
|  | No | 7 (53.8%) |
| **Dead** | Yes | 3 (23.1%) |
|  | No | 10 (76.9%) |
|  |  | **Median**<br>**(interquartile range)** |
| **Age** |  | 4 (3-11) |
| **Infused CD34+ cells (x10<sup>6</sup> cells/kg)** |  | 5.6 (5.3-6.4) |

<sup>a</sup> Cancer types are neuroblastoma<sup>b</sup> and other tumors, including: clear cell renal cell carcinoma<sup>b</sup>, rhabdoid tumor<sup>b</sup>, paraspinal medulloepithelioma, plasmacytoma, Burkitt's lymphoma, embryonal tumor with multilayered rosettes, high-grade neuroepithelial tumor with BCOR (BCL6 corepressor) alteration, acute promyelocytic leukemia, nephroblastoma.
<sup>b</sup> Patients who have relapsed.
G-CSF: granulocyte colony-stimulating factor.

### Sample preparation

Plasma and PBMCs were obtained from whole blood samples of cancer patients. For *in vitro* experiments, PBMCs were obtained from buffy coats and whole blood from 8 healthy donors. PBMCs were obtained by density gradient centrifugation and were cryopreserved in heat-inactivated Fetal Bovine Serum (FBS) (GE Healthcare Hyclone) containing 10% Dimethylsulfoxide (DMSO) (Thermo Scientific Scientific) and stored in liquid nitrogen until they were needed.

Cryopreserved PBMCs were thawed at 37⁰C in a water bath and washed twice with RPMI 1640 medium supplemented with L-Glutamine (Lonza). After that, cells were incubated for 1 hour at 37⁰C and 5% CO_2_ with 10U DNase (Roche) in R10 medium (RPMI 1640 medium containing GlutaMAX from Thermo Fisher Scientific, 10% FBS and 1% Penicillin-Streptomycin (P-S) from Thermo Fisher Scientific). Then, cells were washed once, resuspended in NK cell medium (RPMI 1640 medium containing GlutaMAX, 10% FBS, 1% P-S, 1% MEM Non-Essential Amino Acids Solution and 1% Sodium Pyruvate, both from Thermo Fisher Scientific), filtered using 70µm cell strainers and counted just before further use in flow cytometry experiments or *in vitro* experiments.

### NK cell absolute number determination

The absolute number of NK cells was determined in the whole blood sample. NK cells were identified as CD45+, CD3-, CD56+ and/or CD16+. For that, the following clinical grade fluorochrome conjugated monoclonal antibodies (mAbs) were used: FITC anti-CD16 (CLB/FcGran1), PE anti-CD56 (MY31), PerCP-Cy5.5 anti-CD3 (SK7) and V450 anti-CD45 (2D1) from BD Biosciences. The absolute number of NK cells per μL of blood was calculated based on the following formula: (percentage of NK cells within lymphocyte gate x absolute number of lymphocytes)/100. The absolute number of lymphocytes was obtained from the hemogram.

### DNA extraction, HLA and KIR genotyping

Following manufactureŕs recommendations, DNA was obtained from PBMCs using FlexiGen DNA kit (Qiagen) and. The first step is the addition of lysis buffer to each sample following Qiagen "FlexiGene DNA procedure" flowchart. Briefly, cell nuclei and mitochondria were pelleted by centrifugation and resuspended in denaturation buffer containing QIAGEN Protease. Following protein digestion, DNA was precipitated by addition of isopropanol, recovered by centrifugation, washed in 70% ethanol, and dried. DNA was resuspended in hydration buffer and stored at –20°C for further use.

HLA-A, B and C *loci* typing (necessary to discriminate the KIR ligands present in each patient) was performed using the following kits: LIFECODES HLA-A,B SSO Typing Kit and LIFECODES HLA-C eRES SSO Typing Kit (Lifecodes) all of them CE marked. Lifecodes HLA uses PCR-SSO methodology, subsequent data input on the Luminex® 200™ (Merck) analyzer and results were examined using Matchit DNA software. HLA genotyping was performed at a level of resolution that allowed determining the KIR-binding epitope to distinguish Bw4 specificities and the HLA-C dimorphism at position 80 of the α1 helix. In situations of ambiguity when assigning alleles or to rule out more frequent null alleles, NGS typing was also performed. The NGSgo-MX6 multiplex amplification strategy was used, followed by sequencing on the Illumina Myseq platform and analysis by the NGS engine program.

KIR typing was performed by a PCR-SSP method (sequence-specific primers) using the kit KIR Ready gene kit (Inno-train Diagnostik GmbH). PCR products were amplified and subsequently resolved on agarose gels, and results were interpreted according to the manufacturer’s instructions. Based on KIR gene content, haplotype (A or B) was defined and genotypes (AA or Bx, where “x” can be A or B) were grouped for each patient. The AA genotype is homozygous for inhibitory haplotype A (consisting of 3DL3, 2DL3, 2DP1, 2DL1, 3DP1, 2DL4, 3DL1, 2DS4 and 3DL2). Haplotype B or Bx genotype includes any combination of KIR other than the above.

### NK cell culture with cytokines

For *in vitro* experiments, 1.5x10^6^ PBMCs/mL were cultured in a 48-well plate in 1mL of NK cell culture media (described above) with different combinations of IL-15 (10 ng/mL) and TGF-β (1-10 ng/mL) cytokines. We performed the following conditions: no cytokines (unstimulated), IL-15, 5 ng/mL TGF-β, IL-15 + 1 ng/mL TGF-β, IL-15 + 5 ng/mL TGF-β and IL-15 + 10 ng/mL TGF-β. Cells were cultured for 7 days and media was refreshed at day 4. For the scRNA-seq experiment, PBMCs were cultured in NK cell media without cytokines or with a combination of IL-15 (10ng/mL) and TGF-β (1-10 ng/mL) for 4 days.

### scRNA-seq

Four single cell RNA libraries were constructed using a Chromium controller (10x Genomics, Pleasanton, CA, USA) as per the manufacturer’s instructions (10x Genomics Chromium Single Cell 3’ V3.1 chemistry). A total of 12 cDNA amplification cycles and 14 cycles of library amplification were performed. Sequencing was carried out using a Novaseq 6000 SP Reagent Kit v1.5 (Illumina, San Diego, CA, USA; 100 cycles). FASTQ sequence data were aligned to a custom hg38 (GRCh37, CellRanger reference genome version References-2020-A, build GRCh38.p13) using CellRanger count (v7.2.0).

Gene counts were loaded into R (R-project.org/), v4.3.1) using Seurat (v5.0.1). Analysis was conducted with Seurat and extensive use of dplyr (v1.1.4) and tidyverse (v2.0.0) functionality. For each of the four libraries analyzed, we filtered poor quality cells by requiring each cell to have at least 800 expressed genes (of at least one count), at most 40,000 reads (removing cell doublets), and at most 10% mitochondrial RNA (removing dying cells). We then proceed with the normalization, variable feature identification, scaling of the data and PCA of each of the libraries. We next integrated the libraries with the CCA integration method using the IntegrateLayers function in Seurat. UMAP dimensionality reductions were produced, and clusters were computed with the FindNeighbours function followed by FindClusters (resolution of 0.6), resulting in 18 clusters. In addition, phateR (v1.0.7) package was used to calculate 2-dimensional PHATE embeddings from the normalized data.

For the NK cell population, we further annotated cell type clusters of NK cells (clusters 7 and 9) by sub-setting the Seurat object to these cells only, and then repeating the procedure described above, but with only 20 dimensions for the UMAP and clustering. With this, NK cells were further divided into 9 clusters representing NK1, NK2, NK3 and NK4 cells, using expression patterns and marker gene lists. For each NK cell (sub)type, we compared gene expression levels between cells untreated and treated with the IL-15 + TGF-β, using the FindMarkers function in Seurat on the normalized counts (Seurat “RNA” Assay). UMAP and PHATE cell embeddings were visualized using the function Dimplot in Seurat. Gene expression was visualized using Featureplots and dotplots in Seurat.

### Flow cytometry

For flow cytometry analysis of NK cells, thawed PBMCs were first stained with LIVE/DEAD Fixable Aqua Dead Cell Stain Kit (Invitrogen) reagent to exclude dead cells, following manufactureŕs recommendations. Then, cells were washed with PBS containing 2.5% bovine serum albumin (BSA) (Sigma-Aldrich) before performing surface receptor staining. For that, cells were incubated for 30 minutes on ice, in the dark, with the fluorochrome conjugated mAbs summarized in Supplementary Table 3 (for phenotypical analysis 3 different panels were used, numbered from 1 to 3, and an additional panel for *in vitro* experiments). After staining, cells were then washed again with PBS containing 2.5% BSA and resuspended in PBS. Samples were acquired in a LSR Fortessa X-20 flow cytometer (BD Bioscience).

### Flow cytometry data analysis

Flow cytometry data were analyzed using FlowJo v10.8.1. software. Manual and automated analyses were performed. The following plug-ins were used: DownSample (1.1), UMAP, tSNE and FlowSOM (2.6). Briefly, for the automated analysis, events were first downsampled from the gate of interest (NK cells) across all samples using DownSample plug-in and concatenated. For each donor, NK cells were downsampled to 157 cells in the case of the FlowSOM of figure 2 and 1510 cells in the FlowSOM of figure 6. Then, downsampled populations were concatenated for the analysis. FlowSOM was run using the indicated parameters in each figure.

### Determination of cytokines in plasma

Plasma samples were obtained from blood samples and stored at -80⁰C until they were required. For the measurement of IL-15, PDGF-DD and PDGF-BB plasma levels, the Human IL-15, PDGF-DD or PDGF-BB Quantikine ELISA Kits (R&D Systems) were used respectively, following manufacturer’s recommendations. The optical density was determined using Varioskan Flash fluorimeter (Thermo Fisher Scientific) and the standard curve and non-linear regression and log-log line model was performed in GraphPad Prism software. For the measurement of TGF-β plasma levels, Luminex MILLIPLEX TGF-beta 1 Single Plex MAGNETIC Bead Kit (Merck) was used, following manufacturer’s recommendations. Levels of TGF-β were quantified using Luminex® 200™ (Merck) and analyzed using xPONENT® software. For the determination of GDF-15 and PlGF plasma levels, Elecsys GDF-15 and Elecsys PlGF kits (Roche) were used, following manufacturer’s recommendations. For quantification, the electrochemiluminescence was determined in a cobas e 801 analytical unit (Roche) immunoassay analyzer.

### Statistical analysis and data representation

GraphPad Prism v.9.3.1 was used for graphical representation and statistical analysis. Non-parametric Wilcoxon matched-pairs signed-rank test was used to determine significant differences among groups and Mann Whitney unpaired test was used to determine significant differences among non-paired data. Rout test was used in the case of HLA-DR marker in order to identify outliers in CD56^neg^ population. Correlation plots between different variables were calculated and visualized as correlograms. Spearmańs Rank Correlation coefficient was indicated by square size and heat scale. These analyses were performed with R (version 4.3.1): A language and environment for statistical computing (R Foundation for Statistical Computing, Vienna, Austria).

## RESULTS

### NK cell counts, subset frequencies and maturation status after autoHSCT

NK cells phenotype changes during the immune system reconstitution after autoHSCT ^12,24,25,27^. In this work we have studied the NK cell reconstitution in a cohort of 13 pediatric patients with cancer undergoing autoHSCT using blood samples taken at different time points: before autoHSCT (S1), after reaching leucocyte recovery (>1000 leukocytes/µl, usually around day 12 after autoHSCT, S2) and 30 days (S3), 100 days (S4) and 180 days (S5) after autoHSCT. Patients’ clinical characteristics are described in Table 1. After autoHSCT, absolute number of NK cells did not significantly change, although a reduction was noticed from S1 to S2 followed by a sustained increase until S5 (Figure S1A). The percentage of NK cells within lymphocytes did not change during the reconstitution. To characterize NK cells, we defined them by the expression of NKp80 and CD56, and the negative expression of other cell markers (CD3, CD14, CD19 and CD123), as it is shown in Figure S2. We also defined 3 different NK cell subsets based on the expression of the CD56 marker: CD56^bright^, CD56^dim^ and CD56^neg^ ^35^ and we observed that after autoHSCT, their percentage did not overly fluctuate (Figure S1B).

NK cell differentiation and maturation were studied by analyzing the expression of NKG2A and CD57 receptors, which are expressed at early and late stages of differentiation, respectively ^36^. The frequency of NKG2A+ NK cells increased early after transplantation (from S1 to S2) (Figure S1C). In contrast, the frequency of CD57+ NK cells increased at later time-points (>100 days, S4 and S5) (Figure S1D). Also, the co-expression of NKG2A and CD57 was analyzed after autoHSCT for all NK cell subsets. As expected, the results showed that, not only the most immature CD56^bright^ subset were mostly NKG2A+CD57-, but also that these NKG2A+CD57-cells represent the most frequent subset within the total NK cells in these children (Figure S1E). It is important to note that, in order to identify *bona fide* CD56^bright^ (immature) NK cells, we only selected the CD56^bright^ cells lacking CD57 expression, a marker of terminally differentiated CD56^dim^ (mature) NK cells ^37^. It is well known that IL-15, a cytokine with upregulated expression levels after autoHSCT ^25^, induces an increase in CD56 expression in CD56^dim^ NK cells ^38^, independently of their CD57 expression. Overall, we observed that after autoHSCT there is a gradual decrease in the frequency of immature NK cells (NKG2A+CD57-), while the percentage of terminally differentiated (NKG2A-CD57+) NK cells increased. Nevertheless, the more immature subset (NKG2A+CD57-) is the predominant at every analyzed time point (Figure S1E, S1F).

### NK cells are more activated and express higher levels of cytotoxicity-associated markers early after autoHSCT

It was previously reported that genes associated with activation and cytotoxic functions, such as CD38 and CD26 were differentially expressed in adult MM patients after autoHSCT ^26^. Hence, we decided to study the expression of these and other receptors in NK cell subsets in our pediatric cohort. The activating co-receptor and activation marker CD38 ^39,40^ is practically expressed by all NK cell subsets and its expression increased early after autoHSCT (S2), while from day 30 (S3) onwards, the expression gradually decreased to the initial levels (Figure 1). Similarly, autoHSCT induces a transient increase of NK cells expressing HLA-DR and CD26 (Figure 1), which have been described as functionally more activated cells ^41,42^. On the other hand, and regarding the cytotoxicity-associated marker CD8, ^43^, the transient increase at S2 was only observed in the CD56^bright^ NK cell subset (Figure 1).

**Figure 1.**
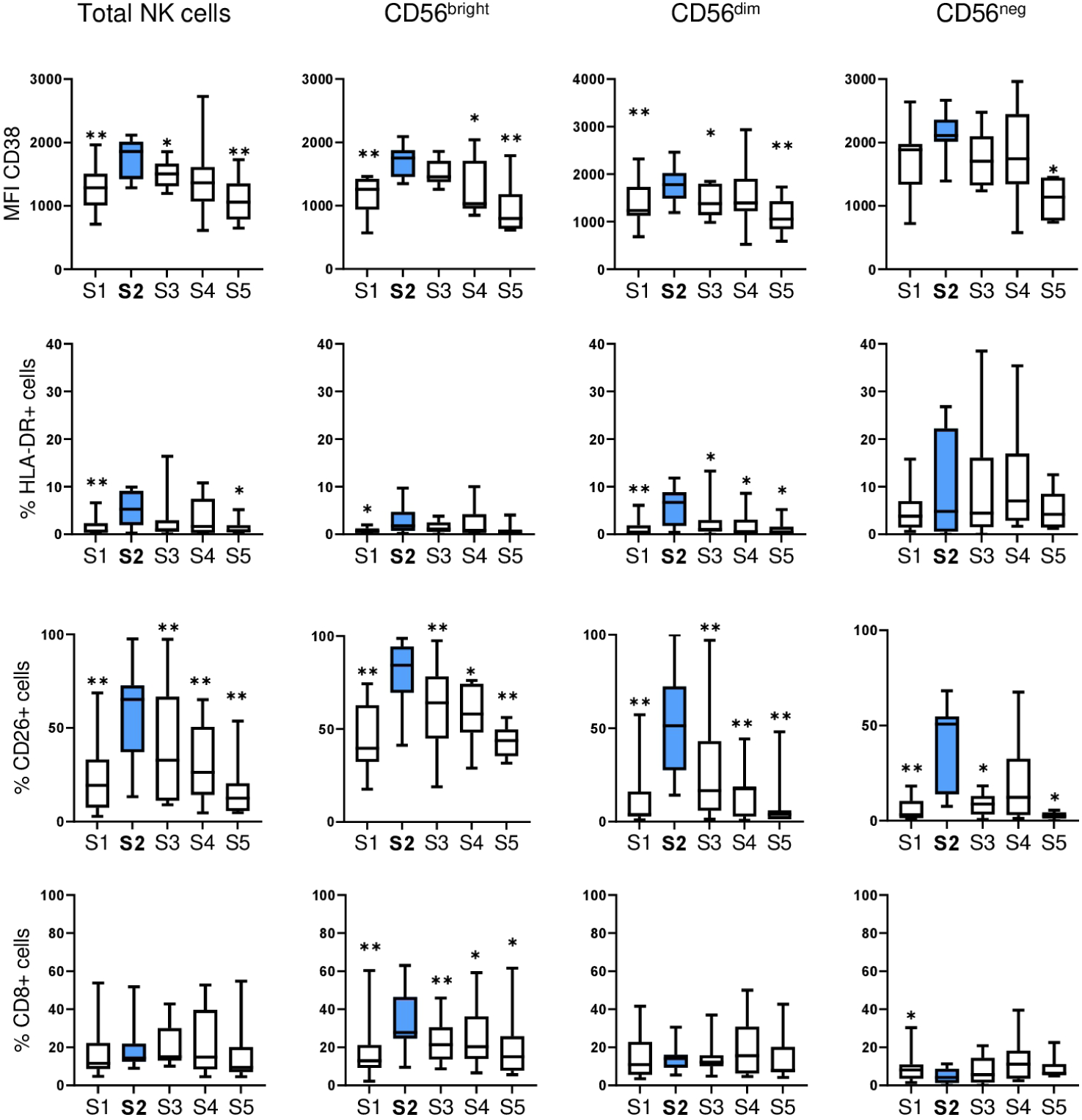
NK cells are more activated early after autoHSCT. Analysis of NK cells in a cohort of pediatric patients before (S1) and following autoHSCT: after reaching leucocyte recovery (more than 1000 leukocytes/µl, usually around day 12 after autoHSCT) (S2), 30 days (S3), 100 days (S4) and 180 days (S5). Boxplot graphs showing the median fluorescence intensity (MFI) of CD38 expression or the percentage of cells expressing the indicated markers in every studied time point within total NK cells and CD56^dim^, CD56^bright^ and CD56^neg^ NK cell subsets. Data are represented as the median and 25–75th percentiles, and the whiskers denote lowest and highest values. A minimum of 10 cells were analyzed in each cell subset. S1 n=12; S2 n=11; S3 n=11; S4 n=9; S5 n=8 for total, CD56^bright^ and CD56^dim^ NK cells. S1 n=11; S2 n=9; S3 n=8; S4 n=9; S5 n=7 for CD56^neg^ NK cells. Results were compared with the S2 time point (blue) using Wilcoxon matched-pairs signed-rank test. *p < 0.05, **p < 0.01.

### NK cell receptor repertoire is altered following autoHSCT

Bulk transcriptomic analyses have shown that the following NK cell receptor genes were differentially expressed in adult patients after autoHSCT: CD31, CD55, CD160 and CD229 ^26^. Hence, we considered that it could be relevant to validate these results in our cohort of pediatric patients by studying the expression of those and other receptors relevant for NK cell function by multiparametric flow cytometry. The results showed that NK cells expressed higher levels of CD31 and LAIR-1 inhibitory receptors at S2 compared to other time points (Figure 2A). CD55, a marker expressed at higher levels on CD56^bright^ NK cells ^44^, followed the same trend (Figure 2A). On the other hand, regarding activating receptors, the percentage of CD160+ NK cells decreased at S2, as well as the expression of CD229 and 2B4 (also known as CD244) (Figure 2B). Lastly, the expression of other receptors, such as Siglec-7, TIM3 and CD226 (DNAM-1) did not significantly change after transplantation (Figure S3A).

**Figure 2.**
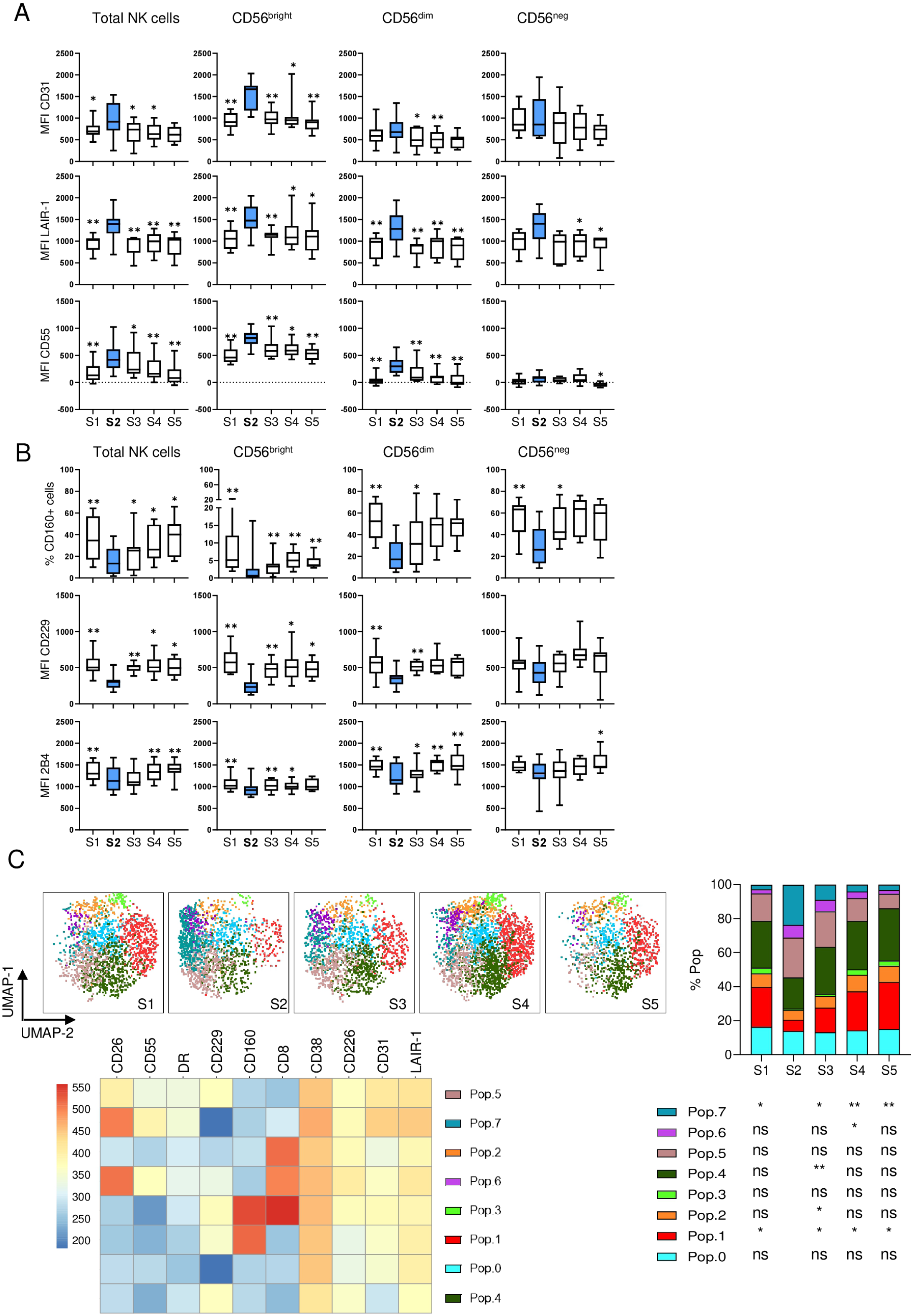
NK cell receptor repertoire is altered early after autoHSCT. Boxplot graphs showing the expression of different receptors in total NK cells and CD56^dim^, CD56^bright^ and CD56^neg^ NK cell subsets at the studied five time points (S1-S5). **(A)** Median fluorescence intensity (MFI) of CD31, LAIR-1 and CD55. **(B)** Percentage of CD160 expressing cells and MFI of CD229 and 2B4 expression. Data are represented as the median and 25–75th percentiles, and the whiskers denote lowest and highest values. A minimum of 10 cells were analyzed in each cell subset. S1 n=12; S2 n=11; S3 n=11; S4 n=9; S5 n=8 for total, CD56^bright^ and CD56^dim^ NK cells. S1 n=11; S2 n=9; S3 n=8; S4 n=9; S5 n=7 for CD56^neg^ NK cells, except for the 2B4 marker (S1 n=12; S2 n=9; S3 n=11; S4 n=9; S5 n=7). **(C)** UMAP projection of CD56+ NK cell populations (Pop) identified by FlowSOM clustering tool for the specified markers, fluorescence intensity of each Pop as indicated in the column-scaled z-score, and bars graph showing the percentage of each Pop at the studied five time points (S1-S5). Results were compared with the S2 time point (blue in A and B) using Wilcoxon matched-pairs signed-rank test. *p < 0.05, **p < 0.01 and ns, no significant.

We also performed an unsupervised analysis using uniform manifold approximation and projection (UMAP) and FlowSOM clustering methodology (Figure 2C). Among CD56+ NK cells, the analysis revealed 8 populations (or Pops) based on the expression of 10 cell surface markers (CD8, CD26, CD31, CD38, CD55, CD160, CD226, CD229, HLA-DR and LAIR-1). We observed a significant increase in the frequency of Pop 7 at S2 corresponding to NK cells expressing high levels of CD26, CD55, HLA-DR, CD38, CD31 and LAIR-1, and low levels of CD229 and CD160. We also observed a significant decrease of Pop 1 at S2 corresponding to NK cells which express high levels of CD229 and CD160, and low levels of CD26, CD55, HLA-DR, CD31 and LAIR-1. Altogether, these results indicate that, early after autoHSCT, NK cells are more activated and that their receptor repertoire is biased towards an inhibitory phenotype, as it was reflected by the transient increase in the expression of inhibitory receptors and the decrease of activating receptors.

To determine whether the previous data are clinically relevant, we performed correlation analyses between the expression of different markers and relapse in the cohort of pediatric patients undergoing an autoHSCT (Figure S3B). We observed a negative correlation between relapse and percentage of CD56^bright^ NK cells early after autoHSCT (S2) and, in accordance with literature regarding adult oncologic patients ^25^, we also detected a positive correlation between relapse and percentage of CD56^dim^ NK cells. This could indicate that patients with a higher frequency of mature NK cells at S2 could have a worse outcome after autoHSCT. On the other hand, CD26 and CD55 expression levels at S2 were negatively associated with relapse. The relevance of these data is very difficult to determine given the number and heterogeneity of the patients. Clearly, we need larger and more homogeneous cohorts of patients to validate these results.

### KIR expression was temporarily increased early after autoHSCT

KIRs regulate NK cell ability to recognize and kill tumor cells and their expression is associated with the outcome of patients with different types of cancer ^30,34,45–48^. We have previously observed that genes codifying different KIRs were overexpressed in adult MM patients early after autoHSCT, such as KIR3DL1, KIR2DL3 and KIR2DS4 ^26^. It has also been described that after alloHSCT there is a sequential recovery of KIRs. KIR3DL1 is the first to recover, followed by KIR2DL2/L3 and finally KIR2DL1 ^49^. Additionally, an upregulation of KIRs expression was noted in CD56^bright^ NK cells early after autoHSCT^24^. Therefore, we decided to study KIR expression in our cohort. Firstly, we analyzed the patientś KIR haplotype (Table S1). Next, we analyzed KIR expression only in the patients who have the genes encoding the corresponding KIRs. We observed an increase in the percentage of KIR3DL1+ cells early after autoHSCT in total NK cells, and an increase of KIR2DL1 and KIR2DL2/L3/S2, only in the CD56^bright^ NK cell subset (Figure S4A-C). However, we did not see significant variations regarding KIR2DS4 after transplant (Figure S4D), a gene expressed only in three patients. On the other hand, no association was observed between KIRs expression and relapse (data not shown).

### Higher KIR expression in mature CD56^dim^NKG2A-CD57+ NK cells and KIR number variation after autoHSCT

During NK cell differentiation, CD56^dim^ NK cells gradually decrease NKG2A and increase CD57 and KIR expression ^36,50,51^. Thus, we next analyzed, within different NK cell subsets, the number of KIR (KIR3DL1, KIR2DL2/L3/S2 and KIR2DL1) that NK cells expressed. First, we observed a significant increase in NK cells expressing one or two KIR early after transplant (S2), followed by a gradual decrease reaching pre-transplant levels at day 180 (S5) (Figure 3A). Regarding CD56^bright^ NK cells, there was a significant decrease in KIR negative cells and an increase in one and two KIR expressing cells at S2 (Figure 3B). Since we have previously seen that the maturation status is altered after autoHSCT (Figure S1E), especially in CD56^dim^ NK cells, we analyzed the frequencies of cells expressing different KIR combinations among different maturation stages. Immature CD56^dim^ NK cells (NKG2A+CD57-) expressed lower KIR levels compared to mature (NKG2A-CD57+) NK cells (Figure 3C). In addition, regardless of the maturation status, CD56^dim^ NK cells had significantly fewer non-expressing KIR cells at S2 (Figure 3C). In conclusion, there is a higher KIR expression in mature terminally differentiated CD56^dim^NKG2A-CD57+ NK cells and higher KIR expression early after autoHSCT at S2 in all NK cell subsets.

**Figure 3.**
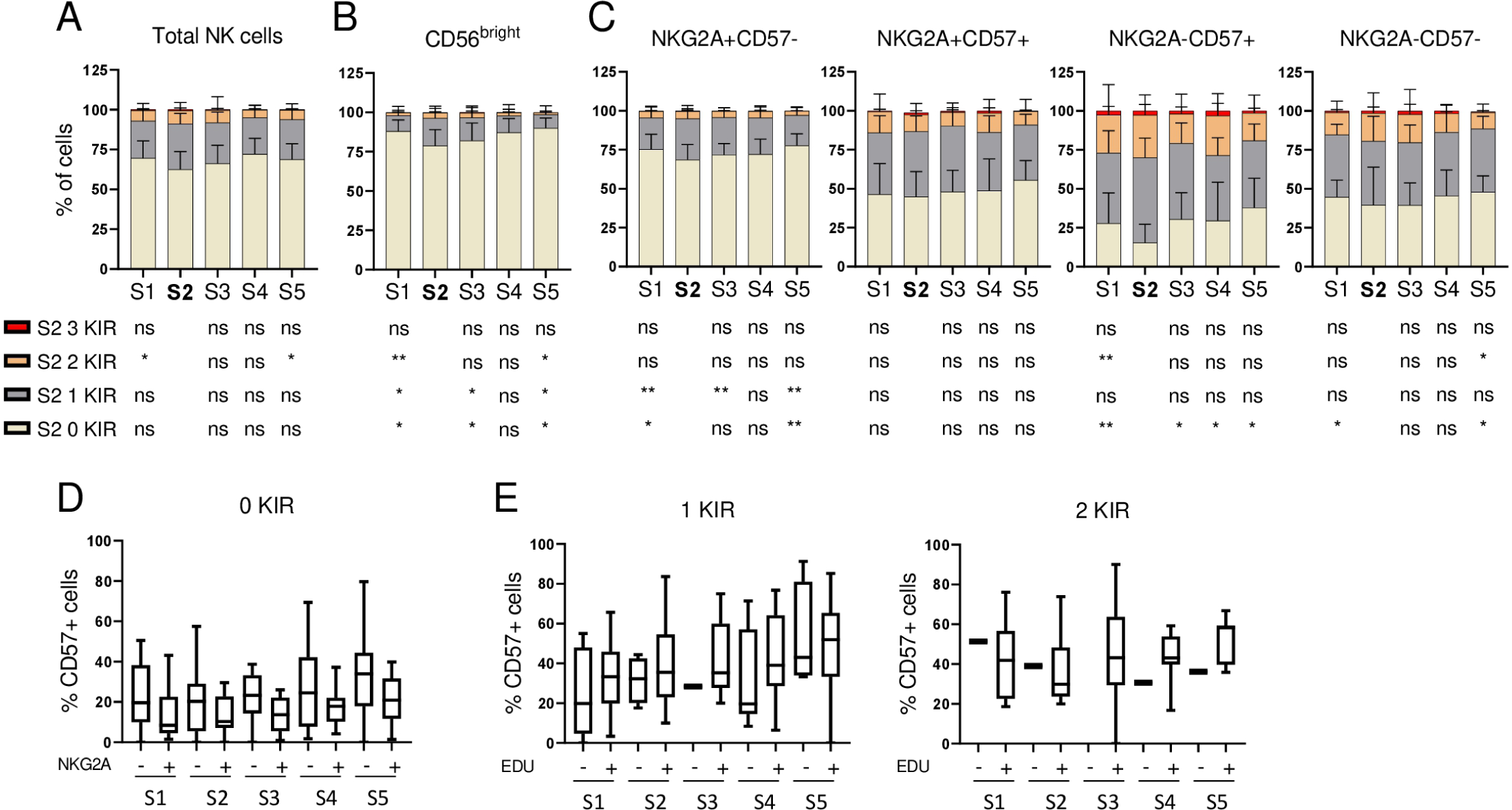
NK cell KIR expression is altered after autoHSCT and NK cell education is not coupled with terminal differentiation. **(A-C)** Bar graphs showing the percentage of cells expressing 0 (light yellow), 1 (grey), 2 (orange) or 3 (red) KIR (KIR2DL1, KIR2DL2/L3/S2 and KIR3DL1) at the studied five time points (S1-S5). Analysis of KIR combinations in total NK cells (A), CD56^bright^ NK cells (B) and CD56^dim^ NK cell subsets based on the expression of NKG2A and CD57 (C). Data are represented as the mean + SD. (S1 n=12; S2 n=11; S3 n=11; S4 n=8; S5 n=8). Results were compared with S2 time point using Wilcoxon matched-pairs signed-rank test. *p < 0.05, **p < 0.01 and ns, no significant. **(D-E)** Boxplot graphs showing the percentage of CD57+ cells at the studied time points on KIR-(0 KIR), 1 KIR or 2 KIR CD56^dim^ NK-cell subsets. Percentage of CD57+ cells in KIR-NKG2A+ or KIR-NKG2A-CD56^dim^ NK cells at different time points (D). Percentage of CD57+ cells in educated (EDU+) or uneducated (EDU-) CD56^dim^ NK cells with 1 or 2 KIR (E). Single KIR positive cells were considered educated if the KIR is specific for the HLA class I haplotype in a given patient (see Table S1). Double KIR positive cells were considered educated if they have at least one KIR that is specific for the HLA class I haplotype in that particular patient (Table S1). Data are represented as the median and 25–75th percentiles, and the whiskers denote lowest and highest values. (S1 n=12; S2 n=11; S3 n=11; S4 n=8; S5 n=8 were analyzed, but only data that followed the education criteria specified above were considered, and more than one value could be added for each patient because NK cells could be educated for more than one KIR). Results were compared between – and + samples for each time point using unpaired Mann Whitney test. No significance was not indicated.

### NK cell education and maturation in autoHSCT

NK cells are functionally modulated by interactions between KIRs and NKG2A with self HLA-I molecules in a process called education ^52–54^. Regarding this, in alloHSCT it has been described that education and maturation are parallel, but uncoupled processes ^36^. As we previously observed a reduction in the frequency of immature (NKG2A+CD57-) NK cells and an increment of mature (NKG2A-CD57+) NK cells later after transplant at S5 (Figure S1E), we determined whether the process of education by KIRs and NKG2A can influence the expression of CD57, a marker of terminal differentiation, in our pediatric cohort. The education status of these subsets was determined by analyzing KIR and HLA genotypes in every patient (Table S2). In this context, NK cells were considered as educated if their KIRs were specific for self HLA-I molecules. In each patient, we determined the number of educated and uneducated CD56^dim^ NK cells expressing 1 or 2 KIR, and we analyzed the expression of the CD57 receptor. In NK cells lacking KIRs, the expression of CD57 was compared between NKG2A+ and NKG2A-cells (Figure 3D). Single KIR positive NK cells were considered educated if the patient expressed the HLA-I ligand for this KIR, and double KIR positive NK cells were considered educated if the patient expressed at least one HLA-I ligand. The results showed no significant differences in CD57 expression between educated and uneducated subsets after autoHSCT for any number of KIRs studied (Figure 3E). We conclude that NK cell education and maturation are uncoupled processes during immune reconstitution following autoHSCT.

### Transient acquisition of a decidual-like phenotype by NK cells early after autoHSCT

Next, we analyzed the expression of decidual NK cell markers in the pediatric cohort during immune reconstitution, since we have previously discovered that genes codifying CD9 and CD151, two receptors associated with decidual NK cells ^55,56^, were overexpressed in adult MM patients after autoHSCT ^26^. We observed a temporary increase in both CD9 and CD151 expressing NK cells at S2, decreasing to pre-transplant levels at S5 (Figure 4A). This sharp increase is clearly evidenced at S2 where the double positive (CD9+CD151+) NK cell subset is 4 times higher than at S1, which is opposed to the small increase observed for CD9+CD151-NK cells (Figure 4B). This result is consistent among all NK cell subsets (CD56^bright^, CD56^dim^ and to a less extent in CD56^neg^). It has been previously shown that CD151 expression is higher in decidual NK cells, followed by circulating CD56^dim^ cells, while circulating CD56^bright^ cells express very low levels of this receptor ^56^. However, in our cohort, we observed that CD151+ cells expressed lower levels of CD56 than CD151-cells (Figure 4C). Additionally, we analyzed CD49a, another decidual NK cell marker ^55^, and we observed low expression levels that, nevertheless, were significantly increased early after autoHSCT (Figure 4D). Altogether, these results suggest that, following autoHSCT, NK cells acquire a decidual-like phenotype supporting previous results at the transcriptomic level.

**Figure 4.**
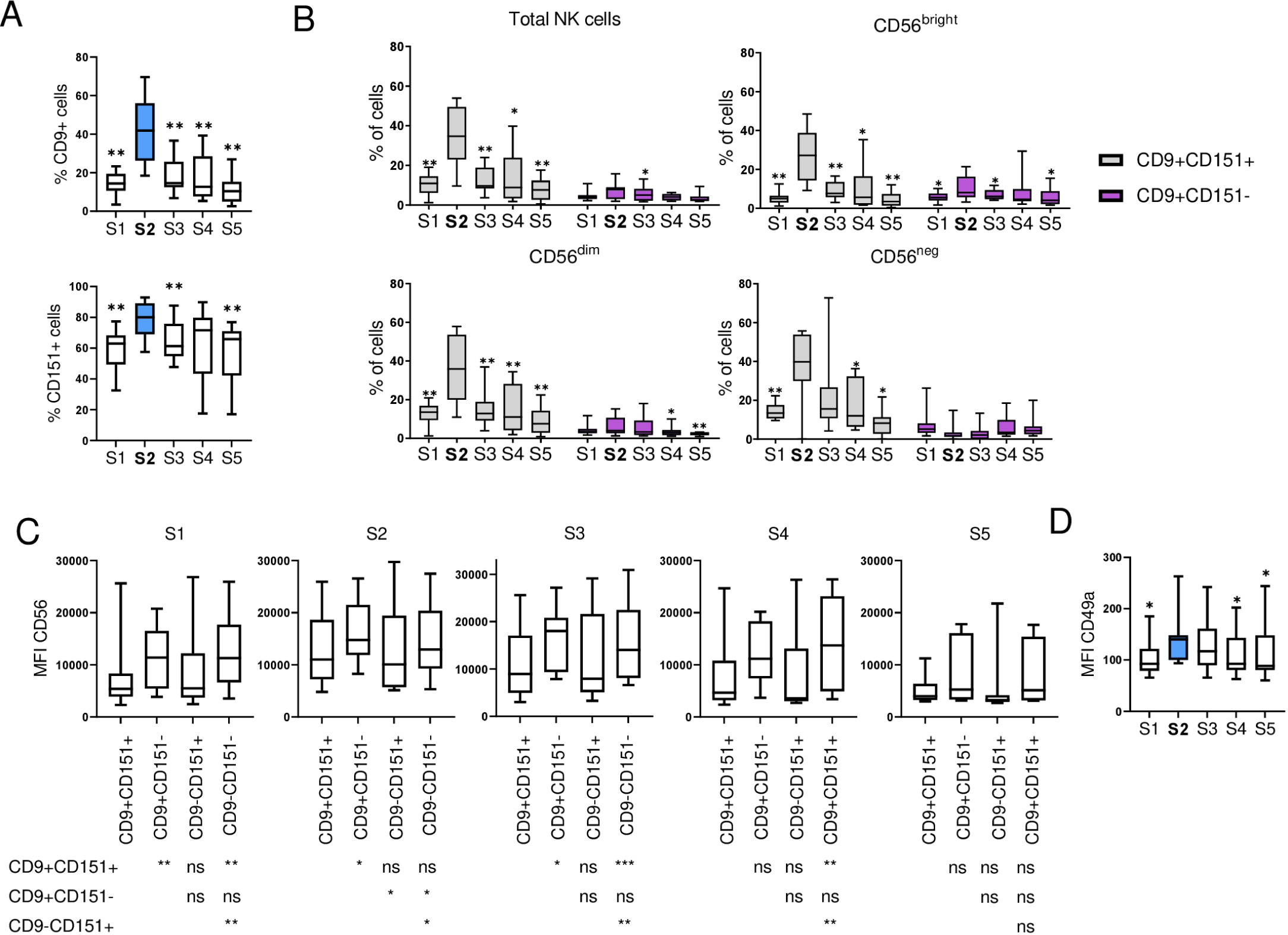
Transient acquisition of decidual-like phenotype by NK cells early after autoHSCT. Boxplot graphs showing the analysis of total NK cells and CD56^dim^, CD56^bright^ and CD56^neg^ NK cell subsets at the studied five time points (S1-S5). **(A)** Percentage of cells expressing CD9 (upper graph) and CD151 (lower graph) in total NK cells. **(B)** Percentage of CD9+CD151+ (grey) and CD9+CD151-(purple) within total NK cells (grey) and in the three NK cell subsets. **(C)** Median fluorescence intensity (MFI) of CD56 in CD9+CD151+, CD9+CD151-, CD9-CD151+ and CD9-CD151-in CD56+ NK cells. **(D)** MFI of CD49a in total NK cells. Data are represented as the median and 25– 75th percentiles, and the whiskers denote lowest and highest values. (S1 n=12; S2 n=11; S3 n=11; S4 n=9; S5 n=8). Results were compared with the S2 time point using Wilcoxon matched-pairs signed-rank test. *p < 0.05, **p < 0.01 and ns, no significant.

### Plasma cytokine profile following autoHSCT

In a previous publication, we have hypothesized that the cytokine *milieu* that is present immediately after autoHSCT may be responsible for the acquisition of a decidual-like phenotype by NK cells ^26^. Therefore, to address this question, we have measured the plasma levels of molecules with important roles in NK cell biology and that may have an impact in the expansion of a decidual-like NK cell subset during autoHSCT.

IL-15 is a cytokine with an important role in proliferation, development, cytotoxic activity and cytokine production of NK cells ^57–59^. Similar to what we and others previously described in adult patients ^9,25^, we found that IL-15 plasma levels also significantly increased early after autoHSCT in our pediatric cohort (Figure 5A). Interestingly, we found a positive correlation between IL-15 levels at S2 and the expression of HLA-DR, CD26, CD55 and CD38 (Figure 5B). These markers also increased at S2, suggesting that increased levels of this cytokine are, at least partially, responsible for the activation phenotype of NK cells at S2.

**Figure 5.**
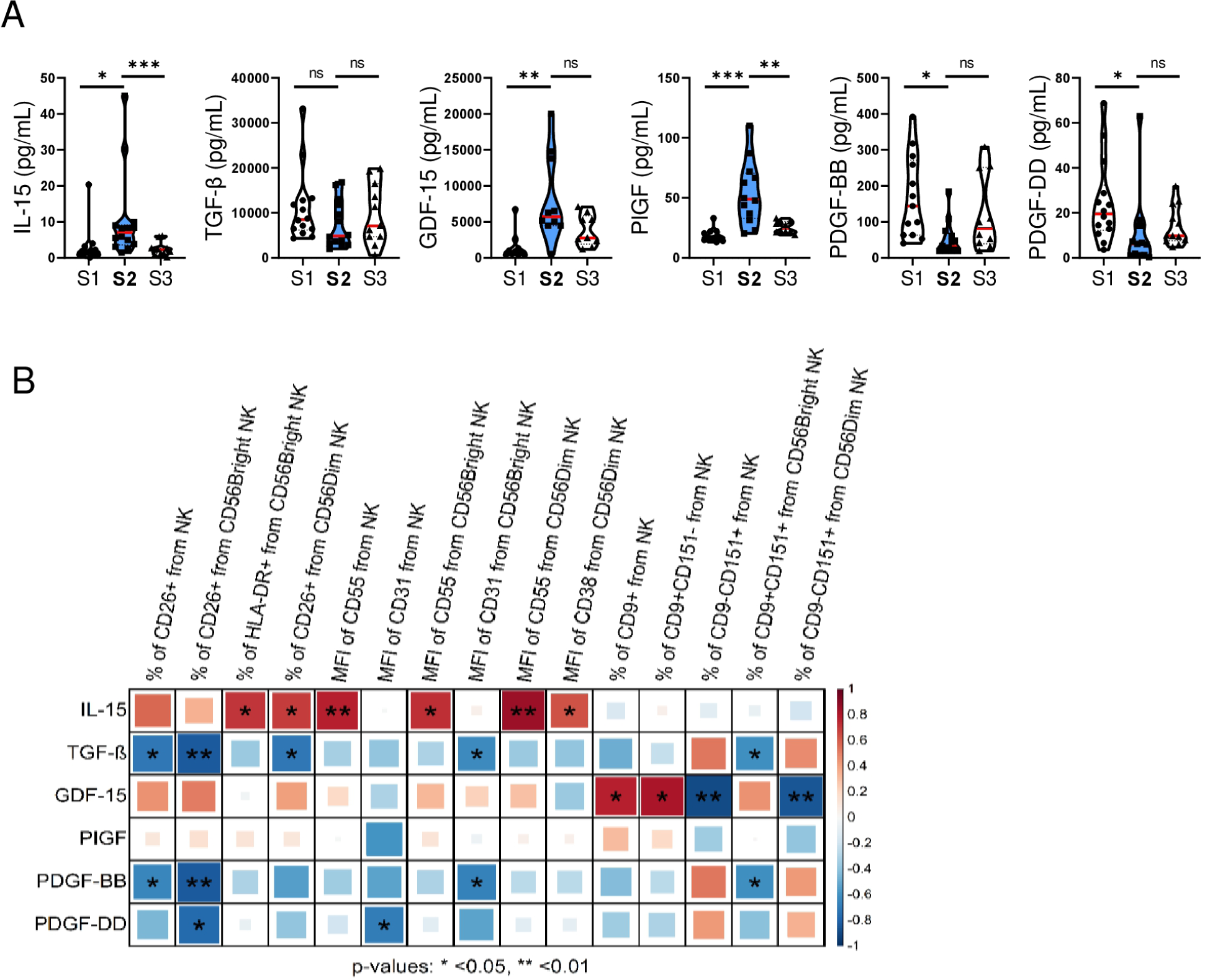
Cytokines plasma levels are altered early after autoHSCT. **(A)** Violin plots showing IL-15, TGF-β, GDF-15, PlGF, PDGF-BB and PDGF-DD plasma levels in a cohort of pediatric patients before (S1) and after autoHSCT: after reaching leucocyte recovery (more than 1000 leukocytes/µl, usually around day 12 after autoHSCT) (S2) and 30 days (S3). Data are represented with the median (red). (S1 n=13; S2 n=13; S3 n=13). Results were compared with the S2 time point (blue) using Wilcoxon matched-pairs signed-rank test. *p < 0.05, **p < 0.01, ***p < 0.001 and ns, no significant. **(B)** Correlogram showing Spearman correlation of the indicated flow cytometry data at S2 and IL-15, TGF-β, GDF-15, PlGF, PDGF-BB and PDGF-DD plasma levels. *p < 0.05, **p < 0.01.

TGF-β has been shown to modulate NK cells receptor repertoire and is produced by tumor, including NBL cells, and healthy cells ^60–62^. This molecule also inhibits NK cell– mediated cytotoxicity and favors immune evasion ^63^. In our cohort, we observed that plasma levels of TGF-β slightly decreased, although it was not statistically significant (Figure 5A). Additionally, a negative correlation was observed at S2 between plasma levels of TGF-β and the increased expression of CD26 and CD31 (Figure 5B). On the other hand, we also analyzed the plasma levels of growth differentiation factor 15 (GDF-15), which has a role in the development of NK cell dysfunction during systemic inflammation ^64^. We found that GDF-15 levels significantly increased at S2 (Figure 5A) and that positively correlated with the frequency of CD9+ NK cells (Figure 5B), a subset that is very elevated at S2.

Placental growth factor (PlGF) is a pleiotropic cytokine which regulates the maturation of uterine NK cells and upregulates the expression of other angiogenic factors such as vascular endothelial growth factor (VEGF) ^65,66^. PlGF levels significantly increased at S2 (Figure 5A). However, no significant association of PlGF levels with CD9+ NK cells, or any other NK cell subset, was observed (Figure 5B). Platelet-derived growth factors (PDGFs) are relevant mitogens for many cell types ^67^ and many tumors use them as an autocrine signal for proliferation ^68–70^. NK cells express receptors for these molecules ^71^ and PDGF-BB has been described to be associated with progression-free survival (PFS) in NHL patients who have undergone autoHSCT ^9^. In our cohort, the plasma levels of PDGF-BB and PDGF-DD significantly decreased early after autoHSCT (Figure 5A). Also, we observed a negative correlation at S2 between these PDGFs and the expression levels of CD31 as well as the frequencies of CD26+ and CD9+CD151+ NK cells (Figure 5B), which were increased at this point. To sum up, early after autoHSCT IL-15, GDF-15 and PlGF levels increase, PDGF-BB and PDGF-DD levels decrease while the concentration of TGF-β in plasma slightly changes.

### IL-15 and TGF-β cooperate to induce a decidual-like phenotype on NK cells

It has been reported an acquisition of decidual features, such as expression of CD9, by peripheral blood NK cells by their culture with TGF-β ^72,73^. Nevertheless, in our cohort the expansion of a decidual-like NK cell subset following autoHSCT cannot be attributed only to TGF-β levels, since we observed no significant changes after transplant (Figure 5A). This prompted us to consider the possibility that a combination of cytokines may be required for the expansion of this specific NK cell subset. In fact, previous publications have shown that the combination of TGF-β with IL-15 induced the acquisition of decidual NK cell markers ^74,75^. Therefore, given that in our pediatric cohort the IL-15 levels significantly increased at S2, while TGF-β levels did not, we performed *in vitro* experiments to determine whether the combination of these two cytokines recapitulates the NK cell phenotype observed after autoHSCT, including the decidual-like subset expansion, at S2.

To study the effect of IL-15 and TGF-β at single-cell resolution, we performed a single-cell RNA sequencing experiment (scRNA-seq) on PBMC cells from a healthy donor cultured with and without the presence of IL-15 (10 ng/mL) + TGF-β (5ng/mL). Principal component analysis (PCA) and UMAP mapped these cells into 18 clusters representing the mayor immune cell populations, including B and T cells, monocytes, NK cells, and others (Figure 5SA). By further sub-clustering NK cells, we identified 4 different NK sub-populations, namely NK1, NK2, NK3 and NK4 (Figure S5B). We followed the nomenclature by Crinier et al. ^76^, and NK1 corresponded to CD56^dim^ and NK2 to CD56^bright^ NK cells. NK1 cells were exclusively present on unstimulated NK cells while NK2 cells were observed in both unstimulated and IL-15 + TGF-β treated NK cells. On the other hand, clusters NK3 and NK4 were almost exclusively observed in IL-15 + TGF-β treated NK cells.

We used a complementary method of dimensionality reduction, namely PHATE (Potential of Heat-diffusion for Affinity-based Transition Embedding), to further visualize the NK cells, and observed that the NK4 cluster was only distantly related to the cluster NK1, suggesting that cytokine stimulation has triggered a transcriptional program that significantly differs from the one observed in unstimulated NK cells (Figure 5SC). Determination of the cell cycle state based on the expression of G2/M and S phase markers, indicated that while the majority of NK4 cells were in the cycling phases (G2/M and S), only 49% and 64% of cells in the NK3 and NK2 clusters, respectively, were actively cycling. Virtually none were observed in the NK1 cluster (Figure 5SD, 5SE and 5SF). Finally, we analyzed the expression levels of genes that encoded cell surface markers that significantly changed early after autoHSCT (S2) in our cohort of pediatric patients. We observed significant upregulation of *DPP4* (CD26), *CD9* (CD9), *ITGA1* (CD49a) and *CD8A* (CD8) in treated cells, and downregulation of *CD160* (CD160) and *CD244* (2B4) (Figure 5SG, 5SH and 5SI), consistent with what has been observed at the protein level at S2. The significant gene expression changes observed in *CD38* and *PECAM1* did not correlated with cell surface expression of CD38 and CD31, respectively, at S2. Genes encoding other cell surface receptors did not significantly changed.

We also isolated PBMCs from several healthy donors (n=7) and incubated them with IL-15 and TGF-β at increasing concentrations. We studied the following conditions: unstimulated, 5 ng/mL of TGF-β, 10 ng/mL of IL-15 without or with increasing concentrations of TGF-β (1, 5 or 10 ng/mL). NK cells were identified by flow cytometry as viable CD3-/CD14-/CD19-/CD123-/CD56+ cells and were analyzed to determine the differentially expressed markers. Conventional supervised analyses were performed and higher CD38, HLA-DR, CD55, CD56 and CD151 expression levels were observed when cells were incubated with IL-15 compared with the unstimulated condition (Figure S6A-D). On the other hand, TGF-β alone induced a significant decrease in the expression of these markers when compared with unstimulated cells, except for CD151 (Figure S6A-D). Nevertheless, TGF-β tended to down-regulate IL-15-induced expression of CD38, HLA-DR, CD55 and CD56, but not to the levels of unstimulated NK cells, suggesting that, at these concentrations, IL-15 has a more dominant effect than TGF-β. On other hand, we observed a higher expression of decidual markers CD9 and CD49a in conditions containing both TGF-β and IL-15, suggesting that both cytokines are necessary in order to obtain high levels of these two markers (Figure S6D). Moreover, we observed a significantly lower expression of CD160 and CD229 in response to TGF-β and IL-15 (Figure S6E). The CD160 decrease was even higher when both cytokines were present, suggesting a synergistic effect, while this synergy was not observed when we looked at CD229 expression levels.

To gain more insight we performed an unsupervised analysis using t-distributed stochastic neighbor embedding (tSNE) and FlowSOM clustering tool for total NK cells (Figure 6). We observed an increase in Pops 0, 3 and 4 when cells were cultured with IL-15 and TGF-β in comparison with the unstimulated condition and single cytokine treatment. These Pops define CD9+CD38+CD56^high^ NK cells that, specifically in Pop 0 and 4, also expressed CD55 and CD151. Pop 4, in particular, exhibited the highest expression of CD49a. Pop 1, with a CD9+CD151-CD56^low^ phenotype, was almost exclusively present when NK cells were cultured only with TGF-β and Pop 2, with a CD9-CD151-CD56^low^ phenotype, is the most abundant in unstimulated or only TGF-β conditions. These results suggest that TGF-β is necessary for the expression of CD9 and that IL-15 is responsible for the increased expression of CD56, CD151, CD49a, CD38 and CD55. Within the CD9-NK cells, we found that Pop 5 and 6 were the most abundant when NK cells were cultured only in the presence of IL-15, and that the addition of TGF-β significantly decrease the frequency of these Pops. Furthermore, in several Pops, a TGF-β concentration-dependent effect was evident: Pops 0 and 3 increased with higher TGF-β concentration. Pop 7 is a very residual population (median of 0.53%) characterized by a very high expression of HLA-DR, which may suggest that these cells could be other ILC subset ^77^. More studies are required to properly identify this very small subset. Altogether, these results suggest that IL-15 and TGF-β induce an activated and decidual-like phenotype on NK cells, which is characterized by higher expression of CD56, CD9, CD49a, CD151, CD55, CD38, and HLA-DR markers. We found these cells to be significantly expanded at S2, what correlates with very high levels of IL-15 and unchanged levels of TGF-β.

**Figure 6.**
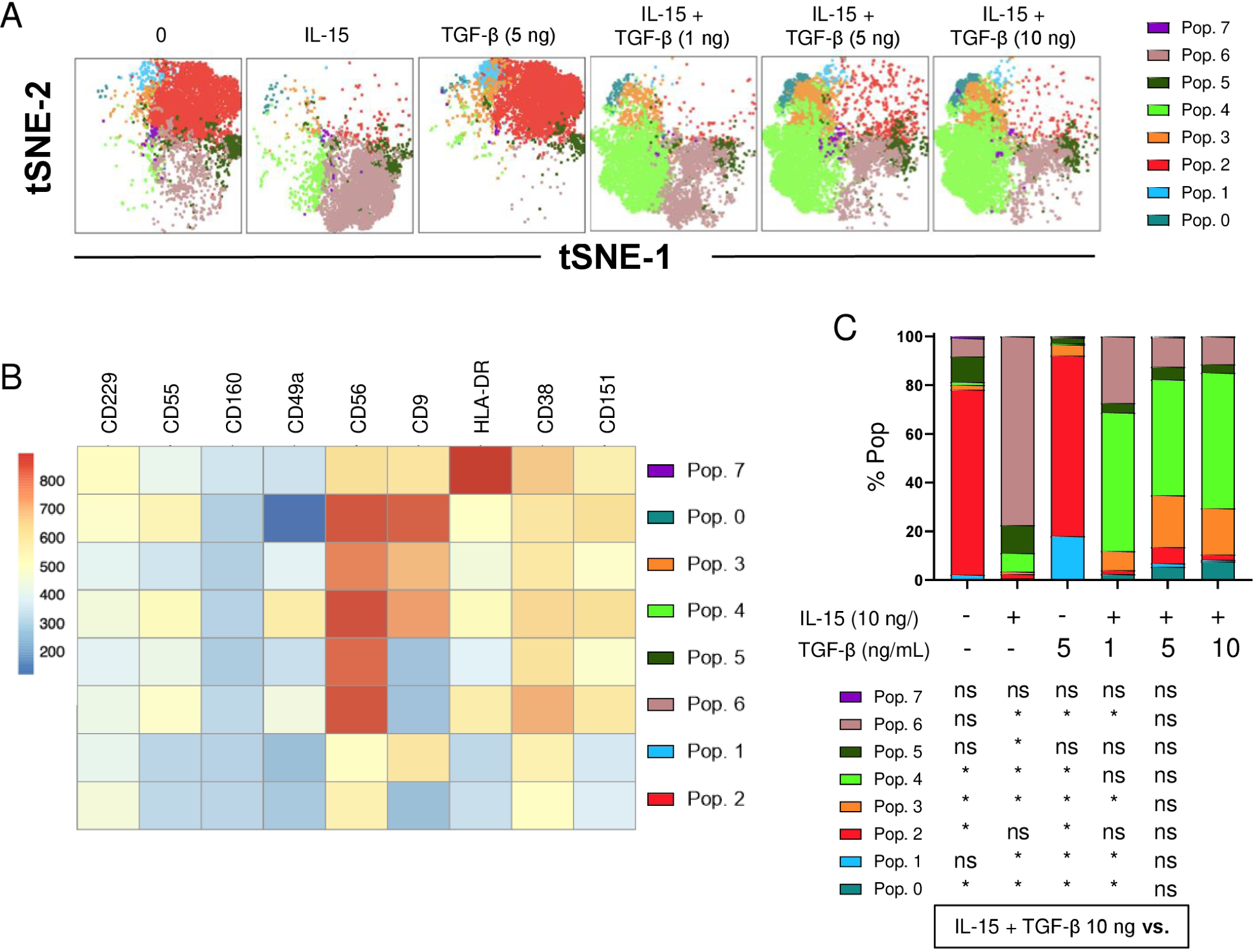
IL-15 and TGF-β induce a NK cell decidual-like phenotype. Unsupervised flow cytometric analysis using tSNE and FlowSOM clustering tools to analyze the indicated markers (CD229, CD55, CD160, CD49a, CD56, CD9, HLA-DR, CD38 and CD151). NK cells were cultured with no cytokines, IL-15, 5 ng/mL TGF-β, IL-15 + 1 ng/mL TGF-β, IL-15 + 5 ng/mL TGF-β, IL-15 + 10 ng/mL TGF-β. IL-15 concentration was 10 ng/mL. (n=7) **(A)** tSNE projection of NK cell populations identified by the FlowSOM clustering tool for the specified markers. **(B)** Fluorescence intensity of each Pop as indicated in the column-scaled z-score. **(C)** Bars graph showing the percentage of each Pop at the studied six conditions. Significance of data was determined using Wilcoxon matched-pairs signed-rank test by comparing each sample with the condition IL-15 + 10 ng/mL TGF-β. *p < 0.05, **p < 0.01 and ns, no significant.

## DISCUSSION

In this work, we have studied the reconstitution of NK cells in pediatric patients with cancer who had undergone autoHSCT. We have analyzed NK cell subsets’ phenotype, plasma levels of cytokines that are relevant in NK cell function, and we have performed *in vitro* experiments (scRNA-seq and flow cytometry) to explore the mechanisms behind the expansion of NK cells with a decidual-like phenotype during immune system reconstitution following autoHSCT. The cell surface phenotype and the transcriptome of NK cells vary considerably after autoHSCT ^24–26^ and, to the best of our knowledge, we have observed a previously undescribed change in NK cells phenotype towards a decidual-like and activated phenotype early after autoHSCT in pediatric patients. This activated and decidual-like phenotype is characterized by an increase of CD56, CD9, CD49a, CD151, CD38, HLA-DR and CD55 expression shortly after autoHSCT. In addition, our *in vitro* experiments suggest that the combination of IL-15 and TGF-β induces, at least partially, this activated and decidual-like phenotype on NK cells.

It is known that TGF-β can induce the acquisition of a decidual-like phenotype in peripheral blood NK cells, characterized, among others, by the expression of CD9 ^72^. However, TGF-β serum levels are reduced after the conditioning regimen for autologous and allogeneic bone marrow transplantation, returning to previous levels 20-50 days post-transplant ^78^. In our study, we did not observe a significant change in TGF-β levels in pediatric patients following autoHSCT (Figure 5A). Other authors have described that the combination of TGF-β with IL-15 induces the acquisition of decidual NK cell markers such as CD9, CD49a and CD151 ^74,75,79^. We and others ^9,25,80^ have shown that IL-15 plasma levels are significantly increased after HSCT (Figure 5A), probably due to transplant-regimen induced depletion of lymphoid cells that consume circulating IL-15. These increased levels of IL-15, together with TGF-β, may contribute to the acquisition of the NK cell decidual-like phenotype. In addition, other cytokines may have a role in the expansion of decidual-like NK cells early after autoHSCT. In fact, we have observed a significant increase in GDF-15 levels, which has been shown to inhibit immune responses, although the mechanism is still unclear ^81^. In addition, there was also an increase in PlGF plasma levels (Figure 5A), which may modulate TGF-β production ^65^. More importantly, we observed that GDF-15 levels associate with CD9 expression at S2 (Figure 5B). Preliminary data from *in vitro* experiments suggest that PlGF and GDF-15, in combination with IL-15 and TGF-β, tend to increase the expression of CD9 and CD49a (data not shown), suggesting that both cytokines could also have a role in the decidual-like phenotype acquisition.

Other authors have shown that NK-92 cells acquire CD9 after coculture with target cells via a trogocytosis-dependent mechanism, leading to a more immunosuppressive cytokine profile and reduced cytotoxic activity ^82^. In agreement with these data, we have previously published that primary CD9+ NK cells exhibited decreased degranulation and lower frequency of MIP-1β+ cells compared to CD9-NK cells in response to PMA and ionomycin ^26^. Nevertheless, we consider that trogocytosis is not a relevant mechanism responsible of CD9+ NK cells expansion in our cohort of patients after autoHSCT. In the autoHSCT scenario, the expansion of CD9+ decidual-like NK cells is transient, occurs early after transplantation, and associates with the increased levels of IL-15 at S2 that, in combination with TGF-β, may be responsible for this phenomenon. However, more studies are needed in order to determine the effect of these and other cytokines in the expression of decidual markers such as CD9.

Several publications have shown an increased frequency of CD9+ circulating NK cells that associates with worse disease evolution ^82–85^ suggesting that CD9 expression on NK cells may be used as a biomarker in cancer patients. Although our study has shown an expansion of CD9+ NK cells, we were not able to find an association between relapse and frequency of these cells. The question that arises is whether the expansion of decidual-like NK cells early after autoHSCT is an event that occurs only in pediatric cancer patients or whether is independent of the underlying condition. We speculate that this event is independent of the malignancy for which patients receive an autoHSCT. In fact, we have previously published that in MM patients there is also an expansion of CD9+ NK cells early after autoHSCT ^26^. Furthermore, it is plausible to assume that this expansion of decidual-like NK cells also occurs in alloHSCT. In this study, we have in depth characterized decidual-like NK cells in pediatric patients and proposed that the different cytokine milieu (i.e. IL-15, TGF-β, etc) during the first days after autoHSCT is responsible for the expansion of this NK cell subset. Evidently, we need larger and more homogeneous cohorts in order to establish CD9+ NK cells as a possible biomarker that may predict the evolution of cancer patients undergoing an autoHSCT in childhood.

On the other hand, the significant decrease in the plasma levels of PDGF-BB and PDGF-DD early after autoHSCT could be due to the lack of signals, such as cytokines, that are required for the secretion of these molecules ^86^. The role of these two growth factors in the expansion and acquisition of the decidual-like phenotype is not known, although our results show that the levels of PDGF-BB inversely correlate with the frequency of CD9+CD151+ NK cells (Figure 5B), suggesting that they may have some role that deserves to be studied.

In addition to the decidual NK cell markers, the cytokine milieu may also have a significant role in the observed changes in the expression of other markers early after autoHSCT. For example, correlation analyses (Figure 5B) and *in vitro* experiments (Figure 6 and S6) showed that the increased expression of CD56, activation markers (HLA-DR and CD38) and receptors such as CD26 and CD55, could be, at least in part, a consequence of the high IL-15 levels at S2 (Figure 5B and S6), as it is shown in other studies ^38,42,87,88^. However, TGF-β tended to counteract the effect of IL-15 (Figure 6C and S6). Remarkably, analysis of data from both scRNA-seq (Figure S5) and flow cytometry (Figure 6 and S6) experiments suggest that the combination of IL-15 and TGF-β has a relevant role in the expansion of decidual-like NK cells early after autoHSCT, at least partially. The two *in vitro* experimental approaches showed that the combination of these two cytokines are responsible for the increased expression of CD9 (*CD9*), CD151 (*CD151*) and CD49a (*ITGA1*), markers that are typical of decidual NK cells ^74,75,79^. Other markers that changed their expression at S2, such as CD26, CD160 and CD229, followed a similar trend of expression after the *in vitro* culture with IL-15 and TGF-β. There were a few markers, especially CD38, in which the scRNA-seq and flow cytometry data somehow were contradictory. While *CD38* mRNA levels decreased following IL-15 and TGF-β treatment (Figure S5), CD38 cell surface expression increased during the culture with these two cytokines (Figure 6 and S6) and at S2 (Figure 1). However, it is important to note that CD38 increased expression was due to IL-15 and that TGF-β decreased it (Figure 6 and S6), and that the higher CD38 expressing Pop6 was the predominant when cells were only treated with IL-15 and they do not express the CD9 decidual marker (Figure 6). In addition, it may be that *CD38* mRNA levels do not correlate with the cell surface expression of this marker. This could happen via differential transcriptional and translational regulations ^89,90^ or via regulation of membrane CD38 internalization ^91^. Undoubtedly, more studies are required to fully understand the role of IL-15 and TGF-β, and very possibly other cytokines, in the acquisition of the unique phenotype that NK cells exhibit early after the autoHSCT.

As expected, the frequencies of CD57+ NK cells in our pediatric patient cohort were lower than those observed in the cohort of MM adults ^25^. This could be owed to the fact that cytomegalovirus infection, more prevalent in adults than in children, induces an expansion of CD57+ NK cells ^92–94^. In addition, we have observed that NK cells express a more immature phenotype early after transplantation at S2 and acquire a more mature phenotype 180 days after autoHSCT, as shown by the increase of the mature NKG2A-CD57+ cell subset at S5. This is in agreement with the model of NK cell development and with the decreased NKG2A and increased CD57 expression during NK cell maturation ^27,36^. This mature NKG2A-CD57+ NK cells exhibited a higher KIR expression and they showed a tendency to expand after autoHSCT at S2 (Figure 3C). These results are in line with reported data from adults receiving an alloHSCT ^36^. Importantly, we have also described for the first time that in autoHSCT, NK cell education and maturation are uncoupled processes, since CD57 expression is not influenced by KIR expression, as it was previously described in alloHSCT ^36,95^.

The search and identification of prognostic biomarkers can be of great help to clinicians. We have observed a positive and significant correlation at S2 between percentage of CD56^dim^ NK cells (Figure S3B), which are more mature than the CD56^bright^ subset, and relapse. Furthermore, CD56^dim^ NK cells encompass a higher frequency of terminally differentiated CD57+ cells than the CD56^bright^ NK cells (Figure S1). Our results are in agreement with data previously published by us and others in which low frequencies of circulating CD57+ and CD57+NKG2A-NK cells are associated with better clinical outcomes ^25,96^. On the other hand, we have also observed that the increased expression levels of CD26 and CD55 at S2 seem to negatively associate with relapse (Figure S3B). It is still unknown how these two receptors are involved in the evolution of autoHSCT and the underlying disease in pediatric patients. CD26 is a dipeptidyl peptidase with a immunoregulatory role in many immune cell types ^97–99^, and some authors have found that high frequencies of transplanted CD26+ lymphocytes have a beneficial effect on NK cell recovery after autoHSCT in MM patients ^100^. Conversely, others have shown that lower CD26 expression on different immune cell subtypes, including NK cells, of the donor stem cell harvest is associated with better survival in alloHSCT ^101^. On the other hand, CD55 is a membrane regulator of C3 activation, and it has been described that, when expressed on NK cells, makes them less effective at killing K562 targets ^102^. Of note, it has been recently published that mouse NK cells presented in the decidua were primarily the differentiated tissue resident NK cell subset, distinguished by their expression of CD55 ^103^. Definitely, it is required to explore if CD55 could be used as a human decidual (and decidual-like) NK cell marker. Nevertheless, these results have to be taken cautiously because our cohort is very small and heterogeneous. To validate these data more studies including a higher number of patients are required.

To the best of our knowledge, this is the first study that has performed such an extended analysis of NK cell subsets in pediatric patients with cancer following autoHSCT. We have demonstrated that there is a shift in NK cells phenotype shortly after autoHSCT. In this setting, NK cells acquire an activated, inhibitory and decidual-like phenotype, and our data suggests that the cytokine milieu plays a role in this process. These changes in NK cell phenotype may have consequences in the autoHSCT evolution and the subjacent cancer. In fact, we have observed a positive correlation between relapse and the percentage of CD56^dim^ NK cells at S2. This could indicate that patients with more mature NK cells at S2 could have a worst prognosis. Although more studies in a larger and more homogenous cohort of pediatric patients are needed to validate these results, our data provides new insights on the biology and physiopathology of NK cells during their reconstitution after autoHSCT, which could help improve the management of these patients and the development of more effective therapies to fight cancer.

## ACKNOWLEDGMENTS

We thank all patients and families who participated in this study, the staff from the Basque Biobank for Research and the staff from the Flow Cytometry and Genetics-Genomics Platforms of Biobizkaia Health Research Institute. We also thank Iñigo Terrén and Rafael González for critical reading of the manuscript.

## FUNDING

This work was funded by the following grants: Fundacion AECC-Spanish Association Against Cancer (PROYE16074BORR) and BIOEF (Basque Foundation for Research and Innovation)-EiTB Maratoia (BIO20/CI/009). GA-P and AA-I are recipients of a predoctoral contract funded by Fundación AECC-Spanish Association Against Cancer (PRDVZ21440ASTA and PRDVZ234209AMAR). DP-A and AT are recipients of a fellowship from the Fundación AECC-Spanish Association Against Cancer (PPLAB212164POLA and PPLAB223829TIJE). AL-P is recipient of a predoctoral contract funded by La Caixa Foundation (100010434; LCF/BQ/DI22/11940012). GA-P and AA-I are recipients of a fellowship from the Jesús de Gangoiti Barrera Foundation (FJGB21/001 and FJBG21/005). BM-M is recipient of a postdoctoral contract funded by CSYF, European Social Fund, Andalucía, Spain (RH-0060-2020). AS is recipient of a grant from the Margarita Salas program, for the requalification of the Spanish university system 2021-2023, financed by European Union -Next Generation EU. LA is an Ikerbasque Research Fellow and FB is an Ikerbasque Research Professor, Ikerbasque, Basque Foundation for Science.

### AUTHOR CONTRIBUTIONS

F.B., G.A.-P. and O.Z. conceived the project; F.B., G.A.-P. and A.O. designed the experiments; G.A.-P., D.P.-A., A.T., M.R., V.S. and L.A. performed experiments; G.A.-P., D.P.-A., S.P.-F., N.G.B. and A.T. conducted data analysis; J.J.U. and I.A. obtained the clinical samples and clinical data from patients; B.M.-M. analyzed HLA and KIR haplotypes; G.A.-P., L.A., V.S., N.G.B and F.B. designed figures; A.O., A.A.-I., A.S., V.S., A.L.-P. and O.Z. provided intellectual input; G.A.-P., F.B. N.G.B. and L.A. wrote the manuscript; all authors critically reviewed the manuscript.

### DATA AVAILABILITY

All data are available in the main text or the Supplementary Materials. scRNA-seq data have been deposited at GEO (Gene Expression Omnibus) under accession number GSE255347.

## Supplemental Information

**This file includes:**

Supplementary Figure 1. NK cell numbers, subsets and maturation status after autoHSCT.

Supplementary Figure 2. Gating strategy for the identification of NK cells.

Supplementary Figure 3. NK cell markers repertoire after autoHSCT.

Supplementary Figure 4. NK cell KIR expression during immune reconstitution after autoHSCT.

Supplementary Figure 5. Transcriptome of NK cells *in vitro* cultured with IL-15 and TGF-β during 4 days.

Supplementary Figure 6. Phenotype of NK cells *in vitro* cultured with IL-15 and TGF-β during 7 days.

Supplementary Table 1. Patientś KIR and HLA haplotypes.

Supplementary Table 2. Education status of NK cells.

Supplementary Table 3. Flow cytometry panels.

---

**Figure S1.**
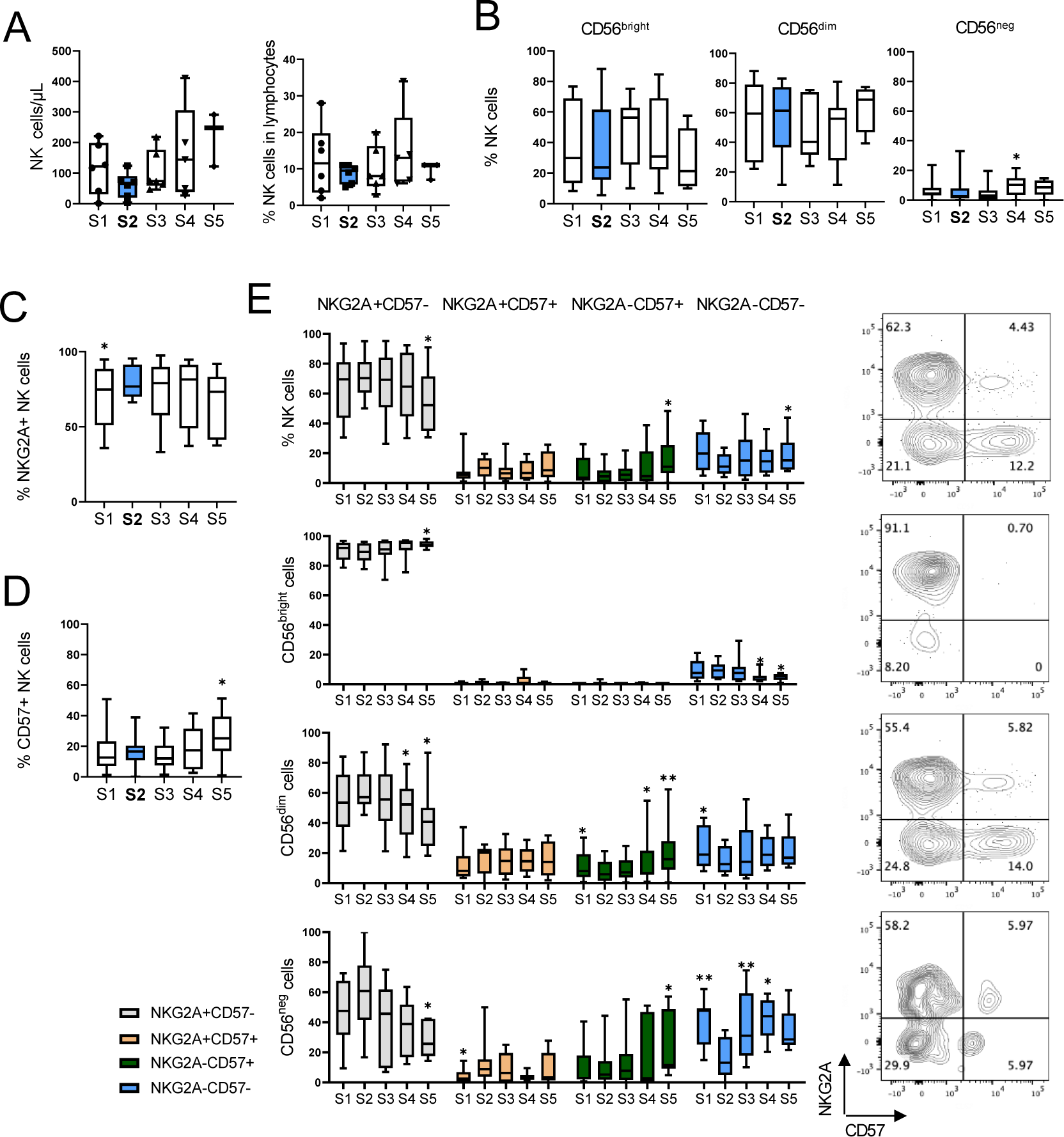
NK cell numbers, subsets and maturation status after autoHSCT. Boxplot graphs showing the analysis of NK cells at the studied five time points: before autoHSCT (S1), after reaching leucocyte recovery (more than 1000 leukocytes/µl, usually around day 12 after autoHSCT) (S2), 30 days (S3), 100 days (S4) and 180 days (S5). **(A)** Absolute number of NK cells (left) and percentage of NK cells within lymphocytes (right) (S1, n=6; S2 n=6; S3 n=6; S4 n=5; S5 n=3). **(B)** Percentage of CD56^bright^, CD56^dim^ and CD56^neg^ NK cell subsets. **(C-E)** Analysis of NKG2A and CD57 receptors expression. **(C)** Percentage of NKG2A expressing NK cells. **(D)** Percentage of CD57 expressing NK cells. **(E)** Percentage of NKG2A+CD57-(grey), NKG2A+CD57+ (yellow), NKG2A-CD57+ (green), NKG2A-CD57-(blue) cells within total NK cells, CD56^bright^, CD56^dim^ and CD56^neg^ NK cells (left) and representative contour plots showing the four populations analyzed (right). Boxplots show the median and 25–75th percentiles, and the whiskers denote lowest and highest values. (S1 n=12; S2 n=11; S3 n=11; S4 n=8; S5 n=8 unless otherwise specified). Significance of data was determined by comparing each sample with sample S2. *p<0.05, **p<0.01 and ns, no significant (Wilcoxon matched-pairs signed-rank test).

**Figure S2.**
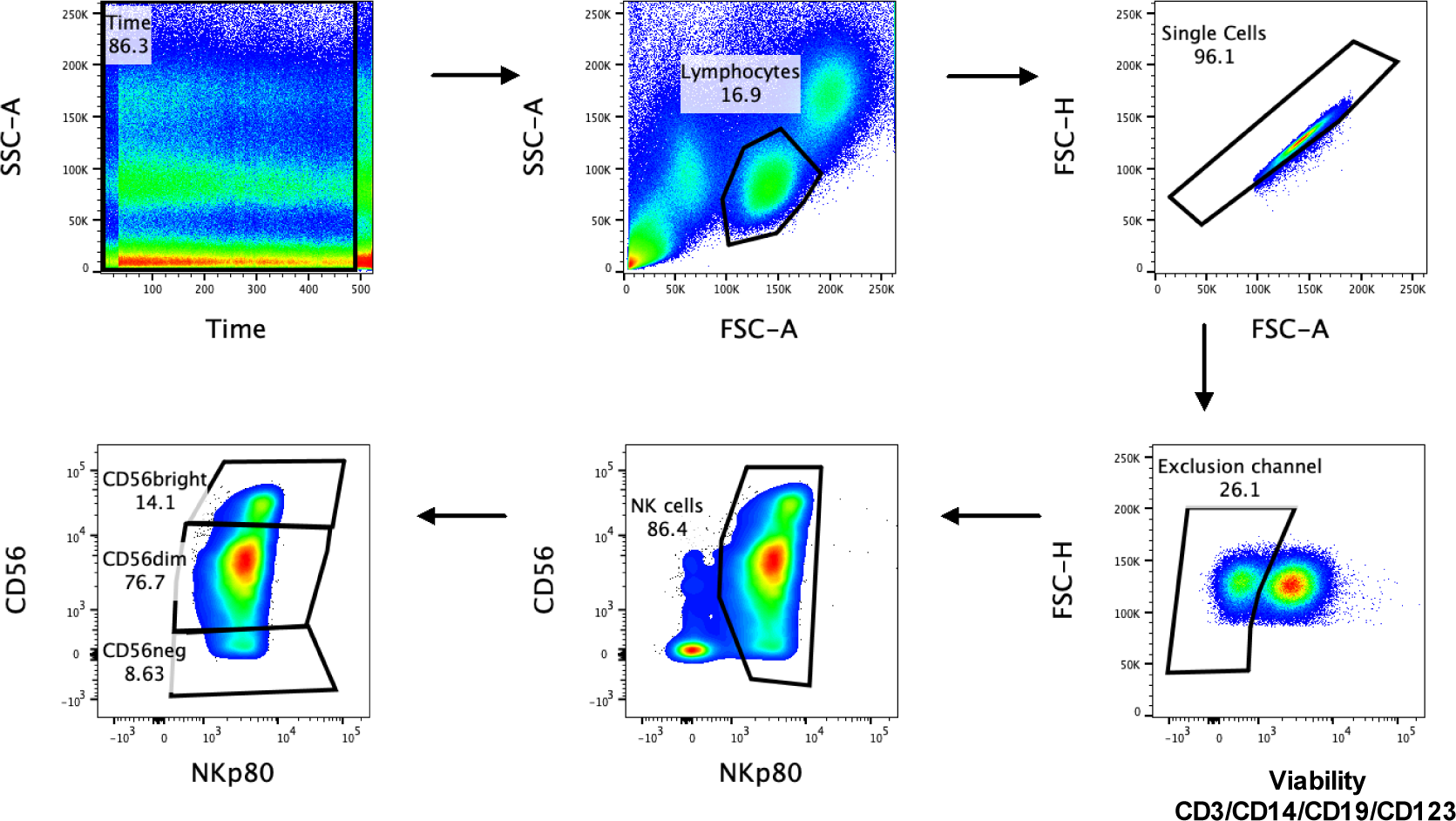
Gating strategy for NK cells identification. Pseudocolor plot graphs representing the gating strategy used for the identification of total NK cells and NK cell subsets. Data from a representative patient is shown. A time gate was made in order to avoid irregular acquisition events, lymphocytes were selected according to their forward and side scatter parameters and, next, single cells were electronically gated. NK cells were identified as negative cells for the exclusion channel (viability, CD3, CD14, CD19 and CD123) and selecting the NKp80+ cells within this population. CD56^bright^, CD56^dim^ and CD56^neg^ NK cell subsets were identified based on the expression of CD56. The expression of different markers were analyzed within total NK cells and NK cell subsets.

**Figure S3.**
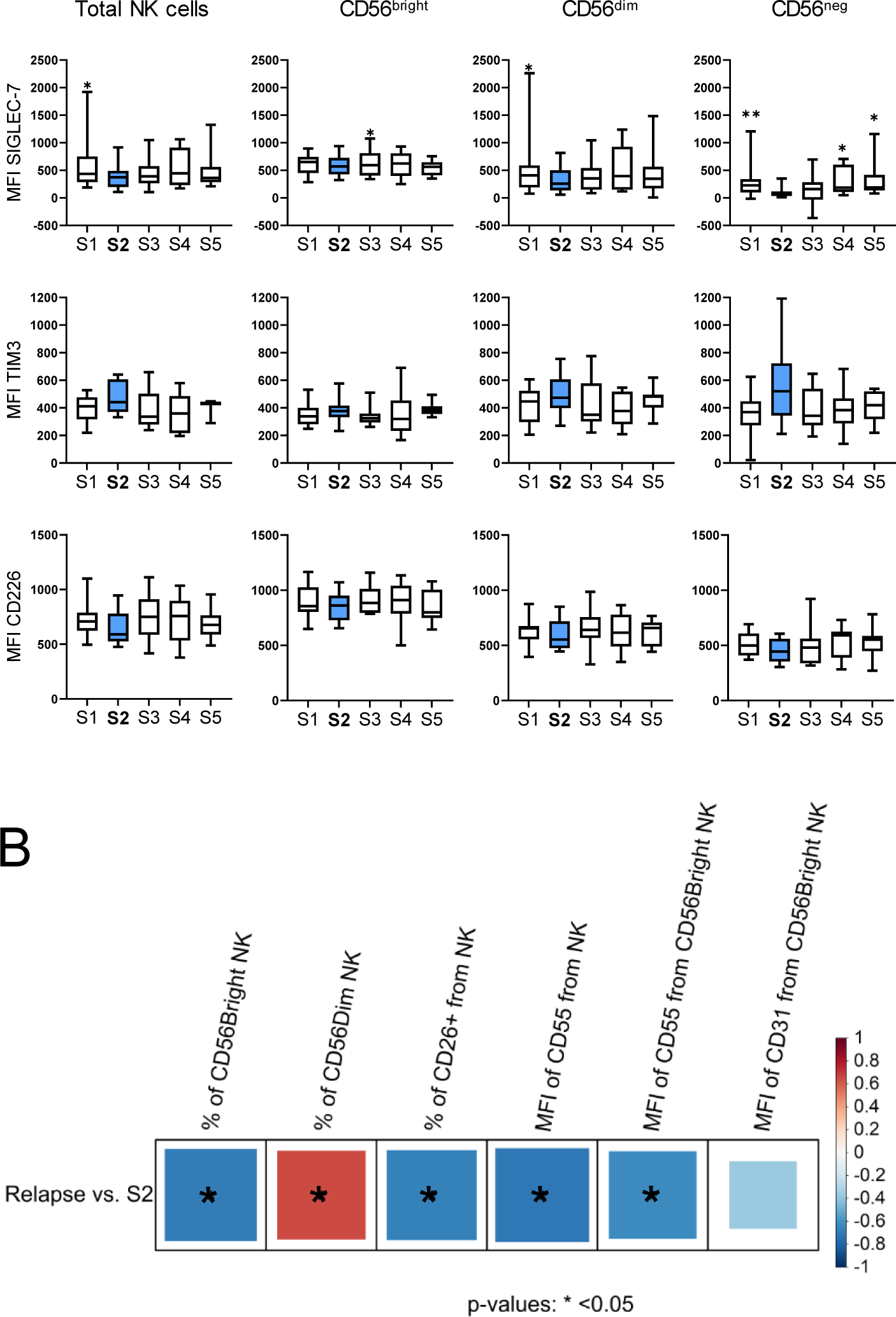
NK cell markers repertoire after autoHSCT. **(A)** Boxplot graphs showing the median fluorescence intensity (MFI) of the specified markers at the studied five time points (S1-S5) within total NK cells and CD56^dim^, CD56^bright^ and CD56^neg^ NK cell subsets. Boxplots show the median and 25–75th percentiles, and the whiskers denote lowest and highest values. (S1 n=12; S2 n=11; S3 n=11; S4 n=9; S5 n=8). Significance of data was determined by comparing each sample with sample S2. *p<0.05, **p<0.01 otherwise, no significant (Wilcoxon matched-pairs signed-rank test). **(B)** Correlogram showing Spearman correlation of the indicated flow cytometry data at S2 and relapse. *p<0.05.

**Figure S4.**
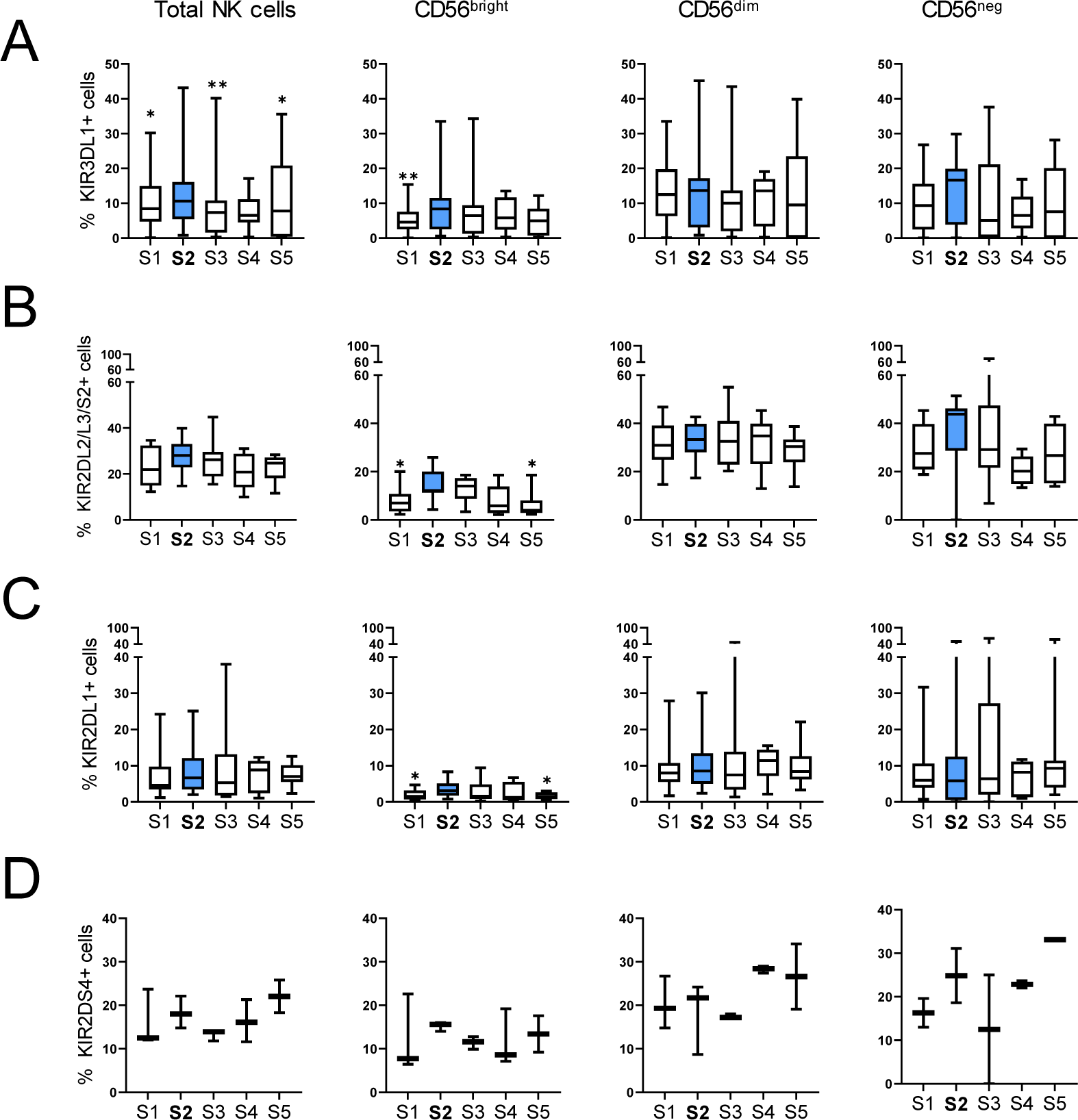
NK cell KIR expression during immune reconstitution after autoHSCT. Boxplot graphs showing, at the studied five time points (S1-S5), the percentage of cells expressing **(A)** KIR3DL1 (S1 n=10; S2 n=9; S3 n=9; S4 n=6; S5 n=6), **(B)** KIR2DL2/L3/S2 (S1 n=12; S2 n=11; S3 n=11; S4 n=8; S5 n=8), **(C)** KIR2DL1 (S1 n=11; S2 n=10; S3 n=10; S4 n=7; S5 n=7) and **(D)** KIR2DS4 (S1 n=3; S2 n=3; S3 n=3; S4 n=3; S5 n=2) within total NK cells and CD56^dim^, CD56^bright^ and CD56^neg^ NK cell subsets. Boxplots show the median and 25–75th percentiles, and the whiskers denote lowest and highest values. Significance of data was determined by comparing each sample with sample S2. *p < 0.05, **p < 0.01 otherwise, no significant (Wilcoxon matched-pairs signed-rank test).

**Figure S5.**
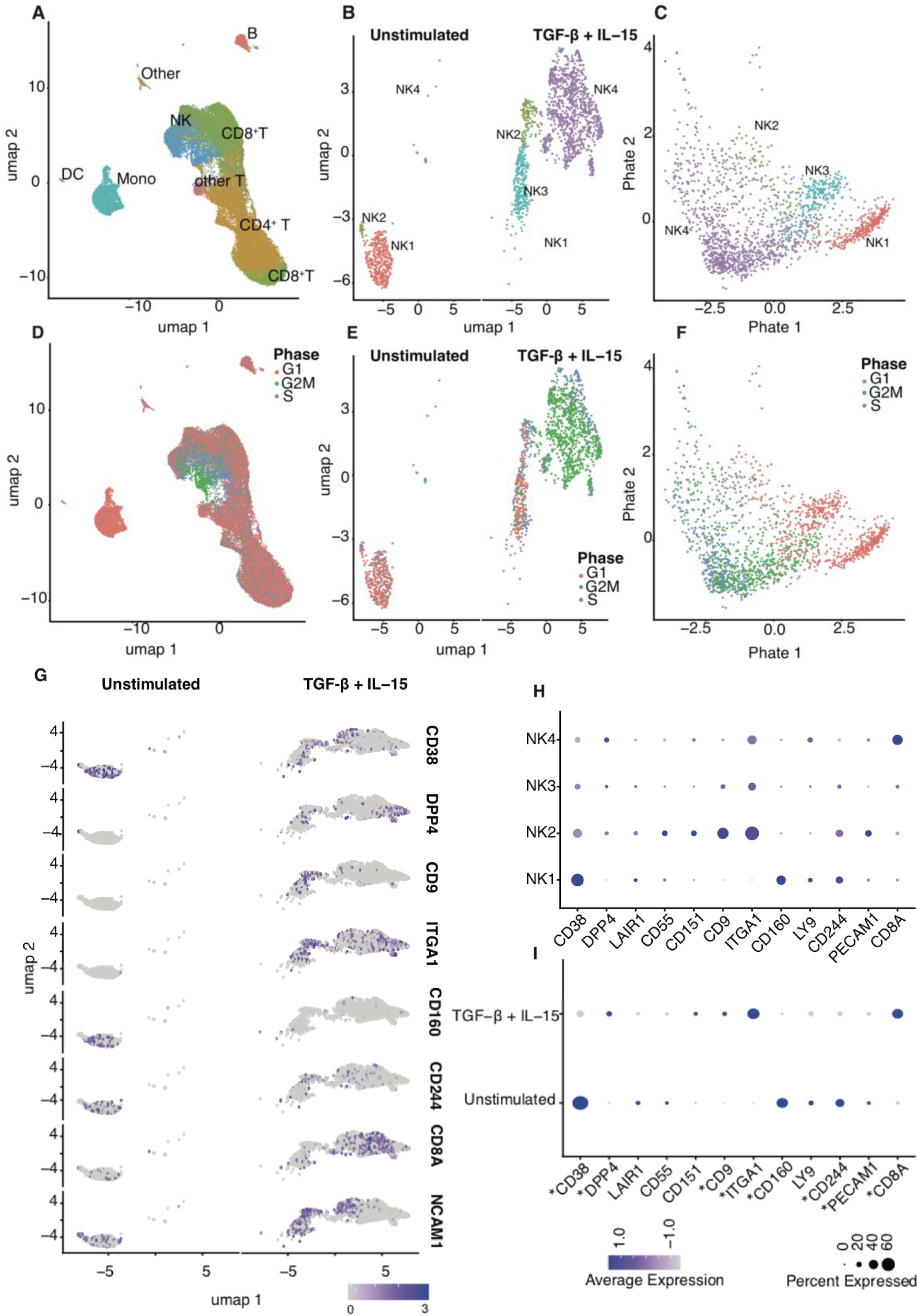
Transcriptome of NK cells *in vitro* cultured with IL-15 and TGF-β during 4 days. scRNA-seq based clustering and cell type identification of 30,842 cells from resting versus stimulated PBMCs shown by an UMAP embedding, colored by the predicted cell type **(A)** and cell cycle phase **(D).** Clustering and cell subtype identification of a subset of 1,708 NK cells (resting and stimulated) as shown by the UMAP **(B, E)** and PHATE embeddings **(C, F),** color coded by cell subtype **(B, C)** and cell cycle phase **(E, F).** UMAP embedding showing expression levels of relevant genes showing differentially significant expression in treated vs unstimulated cells. The panels show resting (left) and stimulated (right) NK cells (**G**). Dot plot of relevant genes. The size of the dots is proportional to the percentage of cells that expresses a given gene within a cell type, while the color of the dot corresponds to the scaled average gene expression **(H, I).**

**Figure S6.**
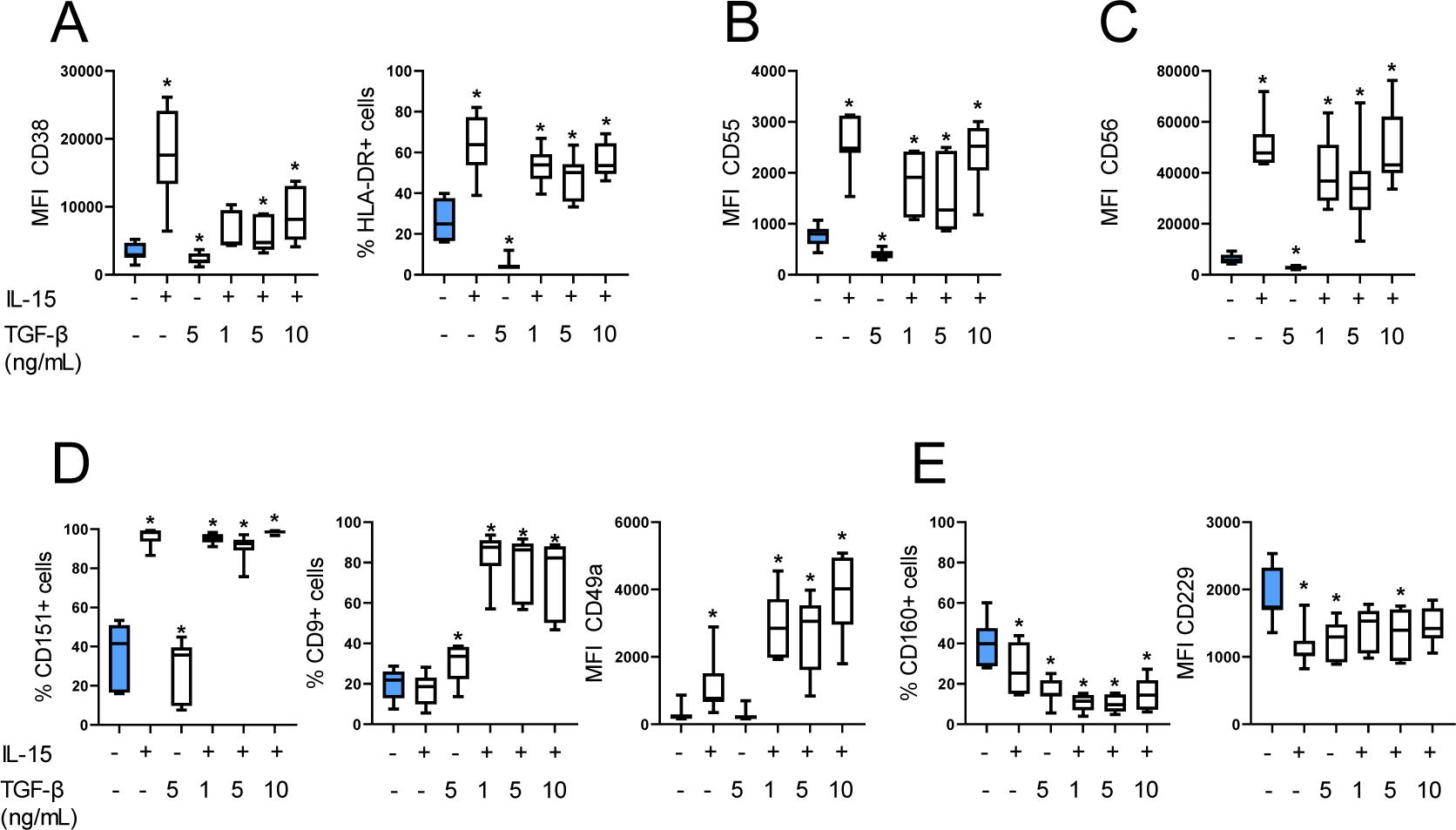
Phenotype of NK cells *in vitro* cultured with IL-15 and TGF-β during 7 days. Boxplot graphs showing the median fluorescence intensity (MFI) or the percentage of cells expressing **(A)** CD38, HLA-DR; **(B)** CD55; **(C)** CD56; **(D)** CD151, CD9, CD49a; **(E)** CD160 and CD229 at the studied six different conditions with IL-15 and TGF-β (unstimulated, 10 ng/mL IL-15, 5 ng/mL TGF-β, IL-15 + 1 ng/mL TGF-β, IL-15 + 5 ng/mL TGF-β, IL-15 + 10 ng/mL TGF-β). PBMCs from 7 healthy individuals were analyzed and NK cells were identified as viable CD3-/CD14-/CD19-/CD123-CD56+ cells. Boxplots show the median and 25–75th percentiles, and the whiskers denote lowest and highest values. Significance of data was determined by comparing each condition with the unstimulated condition (blue) and only significant differences are shown (*p<0.05). Wilcoxon matched-pairs signed-rank test was used.

**Supplementary Table S1.**
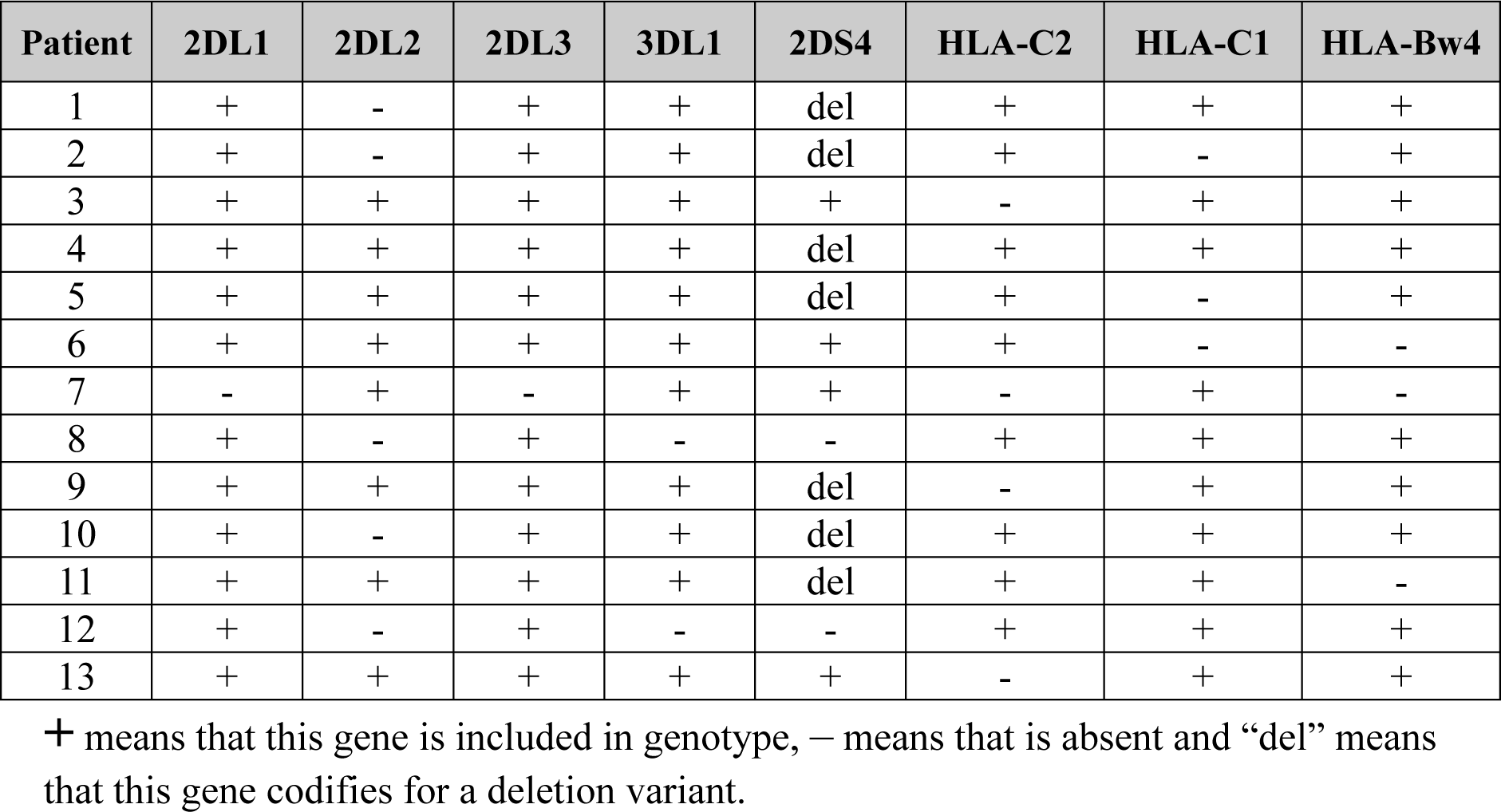
Patientś KIR and HLA haplotypes.

| Patient | 2DL1 | 2DL2 | 2DL3 | 3DL1 | 2DS4 | HLA-C2 | HLA-C1 | HLA-Bw4 |
| --- | --- | --- | --- | --- | --- | --- | --- | --- |
| 1 | + | - | + | + | del | + | + | + |
| 2 | + | - | + | + | del | + | - | + |
| 3 | + | + | + | + | + | - | + | + |
| 4 | + | + | + | + | del | + | + | + |
| 5 | + | + | + | + | del | + | - | + |
| 6 | + | + | + | + | + | + | - | - |
| 7 | - | + | - | + | + | - | + | - |
| 8 | + | - | + | - | - | + | + | + |
| 9 | + | + | + | + | del | - | + | + |
| 10 | + | - | + | + | del | + | + | + |
| 11 | + | + | + | + | del | + | + | - |
| 12 | + | - | + | - | - | + | + | + |
| 13 | + | + | + | + | + | - | + | + |

means that this gene is included in genotype, - means that is absent and “del” means that this gene codifies for a deletion variant.

**Supplementary Table S2.** Education status of NK cells.

| **Patient** | **KIR2DL1**<br>**HLA-C2** | **KIR2DL2/L3**<br>**HLA-C1** | **KIR3DL1**<br>**HLA-Bw4** |
| --- | --- | --- | --- |
| 1 | + | + | + |
| 2 | + | - | + |
| 3 | - | + | + |
| 4 | + | + | + |
| 5 | + | - | + |
| 6 | + | - | - |
| 7 | - | + | - |
| 8 | + | + | - |
| 9 | - | + | + |
| 10 | + | + | + |
| 11 | + | + | - |
| 12 | + | + | - |
| 13 | - | + | + |

+ means educated and - uneducated NK cells for each KIR.

**Supplementary Table S3.**
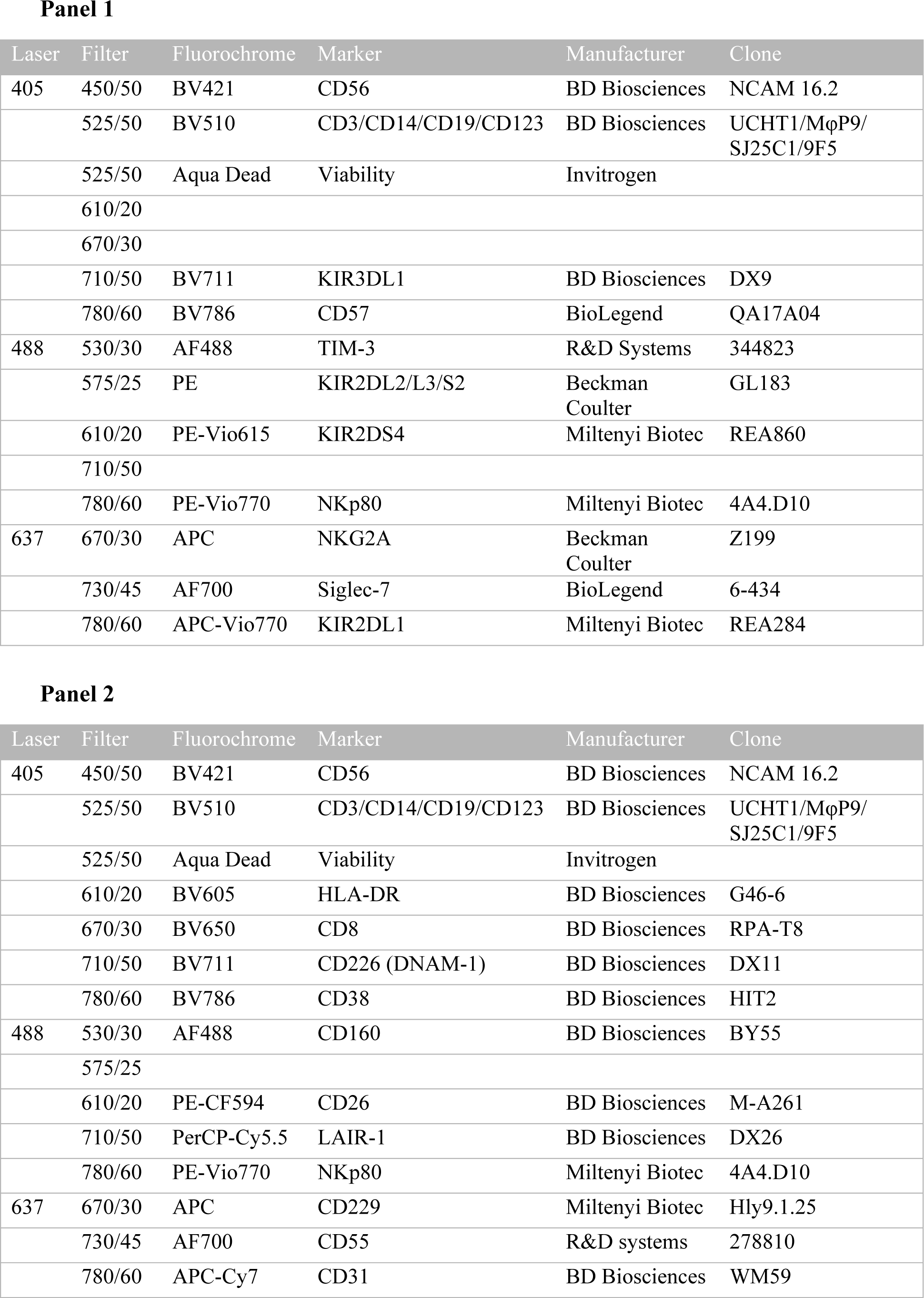
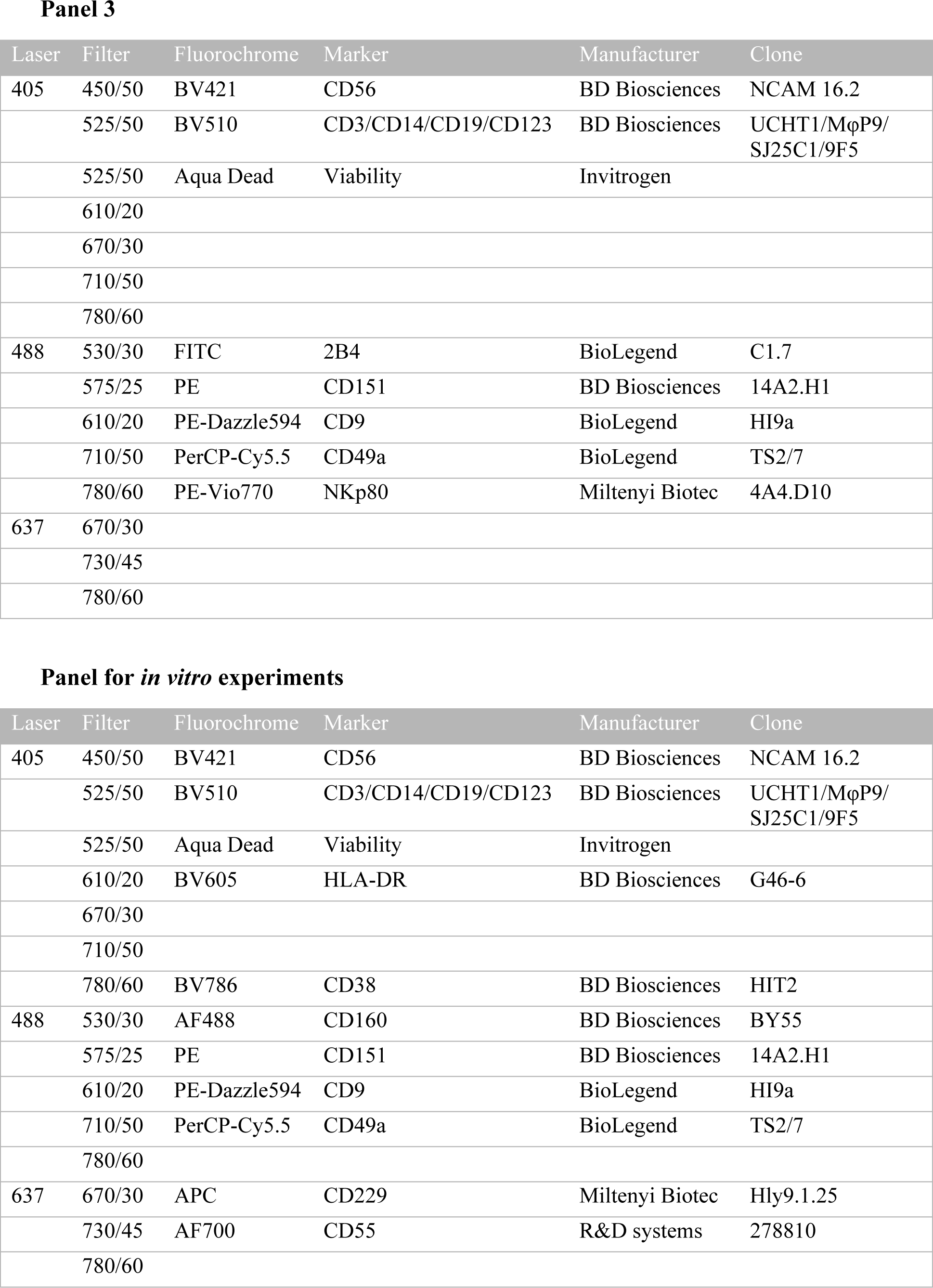
Flow cytometry panels.

## REFERENCES

1 Pinto NR, Applebaum MA, Volchenboum SL, Matthay KK, London WB, Ambros PF et al. Advances in risk classification and treatment strategies for neuroblastoma. J Clin Oncol 2015; 33: 3008–3017.

2 Saleh K, Michot J-M, Camara-Clayette V, Vassetsky Y, Ribrag V. Burkitt and Burkitt-Like Lymphomas: a Systematic Review. Curr Oncol Rep 2020; 22: 33.

3 Passweg JR, Baldomero H, Chabannon C, Basak GW, Corbacioglu S, Duarte R et al. The EBMT activity survey on hematopoietic-cell transplantation and cellular therapy 2018: CAR-T’s come into focus. Bone Marrow Transplant 2020; 55: 1604–1613.

4 Porrata LF, Ingle JN, Litzow MR, Geyer S, Markovic SN. Prolonged survival associated with early lymphocyte recovery after autologous hematopoietic stem cell transplantation for patients with metastatic breast cancer. Bone Marrow Transplant 2001; 28: 865–871.

5 Porrata LF, Gertz MA, Inwards DJ, Litzow MR, Lacy MQ, Tefferi A et al. Early lymphocyte recovery predicts superior survival after autologous hematopoietic stem cell transplantation in multiple myeloma or non-Hodgkin lymphoma. Blood 2001; 98: 579–585.

6 Porrata LF, Inwards DJ, Micallef IN, Ansell SM, Geyer SM, Markovic SN. Early lymphocyte recovery post-autologous haematopoietic stem cell transplantation is associated with better survival in Hodgkin’s disease. Br J Haematol 2002; 117: 629–633.

7 Porrata LF, Inwards DJ, Ansell SM, Micallef IN, Johnston PB, Gastineau DA et al. Early Lymphocyte Recovery Predicts Superior Survival after Autologous Stem Cell Transplantation in Non-Hodgkin Lymphoma: A Prospective Study. Biol Blood Marrow Transplant 2008; 14: 807–816.

8 Rueff J, Medinger M, Heim D, Passweg J, Stern M. Lymphocyte subset recovery and outcome after autologous hematopoietic stem cell transplantation for plasma cell myeloma. Biol Blood Marrow Transplant 2014; 20: 896–899.

9 Porrata LF, Inwards DJ, Micallef IN, Johnston PB, Ansell SM, Hogan WJ et al. Interleukin-15 affects patient survival through natural killer cell recovery after autologous hematopoietic stem cell transplantation for non-Hodgkin lymphomas. Clin Dev Immunol 2010; 2010: 914945.

10 Storek J, Geddes M, Khan F, Huard B, Helg C, Chalandon Y et al. Reconstitution of the immune system after hematopoietic stem cell transplantation in humans. Semin Immunopathol 2008; 30: 425–437.

11 Porrata LF, Litzow MR, Markovic SN. Immune reconstitution after autologous hematopoietic stem cell transplantation. Mayo Clin Proc 2001; 76: 407–412.

12 Porrata LF. Natural Killer Cells Are Key Host Immune Effector Cells Affecting Survival in Autologous Peripheral Blood Hematopoietic Stem Cell Transplantation. Cells 2022; 11: 3469.

13 Vivier E, Artis D, Colonna M, Diefenbach A, Di Santo JP, Eberl G et al. Innate Lymphoid Cells: 10 Years On. Cell 2018; 174: 1054–1066.

14 Guillerey C. Roles of cytotoxic and helper innate lymphoid cells in cancer. Mamm. Genome. 2018; 29: 777–789.

15 Krabbendam L, Bernink JH, Spits H. Innate lymphoid cells: from helper to killer. Curr Opin Immunol 2021; 68: 28–33.

16 Prager I, Watzl C. Mechanisms of natural killer cell-mediated cellular cytotoxicity. J Leukoc Biol 2019; 105: 1319–1329.

17 Michel T, Poli A, Cuapio A, Briquemont B, Iserentant G, Ollert M et al. Human CD56bright NK Cells: An Update. J Immunol 2016; 196: 2923–2931.

18 Cooper MA, Fehniger TA, Caligiuri MA. The biology of human natural killer-cell subsets. Trends Immunol 2001; 22: 633–640.

19 Caligiuri MA. Human natural killer cells. Blood 2008; 112: 461–469.

20 Freud AG, Mundy-Bosse BL, Yu J, Caligiuri MA. The Broad Spectrum of Human Natural Killer Cell Diversity. Immunity 2017; 47: 820–833.

21 Long EO, Sik Kim H, Liu D, Peterson ME, Rajagopalan S. Controlling natural killer cell responses: Integration of signals for activation and inhibition. Annu Rev Immunol 2013; 31: 227–258.

22 Vivier E, Ugolini S, Blaise D, Chabannon C, Brossay L. Targeting natural killer cells and natural killer T cells in cancer. Nat Rev Immunol 2012; 12: 239–252.

23 Cerwenka A, Lanier LL. Natural killer cell memory in infection, inflammation and cancer. Nat Rev Immunol 2016; 16: 112–123.

24 Jacobs B, Tognarelli S, Poller K, Bader P, Mackensen A, Ullrich E. NK cell subgroups, phenotype, and functions after autologous stem cell transplantation. Front Immunol 2015; 6: 583.

25 Orrantia A, Terrén I, Astarloa-Pando G, González C, Uranga A, Mateos-Mazón JJ et al. NK Cell Reconstitution After Autologous Hematopoietic Stem Cell Transplantation: Association Between NK Cell Maturation Stage and Outcome in Multiple Myeloma. Front Immunol 2021; 12: 748207.

26 Orrantia A, Vázquez-De Luis E, Astarloa-Pando G, Terrén I, Amarilla-Irusta A, Polanco-Alonso D et al. In vivo expansion of a CD9+ decidual-like NK cell subset following autologous hematopoietic stem cell transplantation. iScience 2022; 25: 105235.

27 Orrantia A, Terrén I, Astarloa-Pando G, Zenarruzabeitia O, Borrego F. Human NK Cells in Autologous Hematopoietic Stem Cell Transplantation for Cancer Treatment. Cancers (Basel) 2021; 13: 1589.

28 Arteche-López A, Kreutzman A, Alegre A, Sanz Martín P, Aguado B, González-Pardo M et al. Multiple myeloma patients in long-term complete response after autologous stem cell transplantation express a particular immune signature with potential prognostic implication. Bone Marrow Transplant 2017; 52: 832–838.

29 Marra J, Greene J, Hwang J, Du J, Damon L, Martin T et al. KIR and HLA Genotypes Predictive of Low-Affinity Interactions Are Associated with Lower Relapse in Autologous Hematopoietic Cell Transplantation for Acute Myeloid Leukemia. J Immunol 2015; 194: 4222–4230.

30 Venstrom JM, Zheng J, Noor N, Danis KE, Yeh AW, Cheung IY et al. KIR and HLA genotypes are associated with disease progression and survival following autologous hematopoietic stem cell transplantation for high-risk neuroblastoma. Clin Cancer Res 2009; 15: 7330–7334.

31 Leung W, Handgretinger R, Iyengar R, Turner V, Holladay MS, Hale GA. Inhibitory KIR-HLA receptor-ligand mismatch in autologous haematopoietic stem cell transplantation for solid tumour and lymphoma. Br J Cancer 2007; 97: 539– 542.

32 Stringaris K, Barrett AJ. The importance of natural killer cell killer immunoglobulin-like receptor-mismatch in transplant outcomes. Curr Opin Hematol 2017; 24: 489–495.

33 Gabriel IH, Sergeant R, Szydlo R, Apperley JF, DeLavallade H, Alsuliman A et al. Interaction between KIR3DS1 and HLA-Bw4 predicts for progression-free survival after autologous stem cell transplantation in patients with multiple myeloma. Blood 2010; 116: 2033–2039.

34 Erbe AK, Wang W, Carmichael L, Kim KM, Mendoņca EA, Song Y et al. Neuroblastoma patients’ KIR and KIR-ligand genotypes influence clinical outcome for dinutuximab-based immunotherapy: A report from the children’s oncology group. Clin Cancer Res 2018; 24: 189–196.

35 Orrantia A, Terrén I, Izquierdo-Lafuente A, Alonso-Cabrera JA, Sandá V, Vitallé J, et al. A NKp80-Based Identification Strategy Reveals that CD56neg NK Cells Are Not Completely Dysfunctional in Health and Disease. iScience 2020; 23: 101298.

36 Björkström NK, Riese P, Heuts F, Andersson S, Fauriat C, Ivarsson MA et al. Expression patterns of NKG2A, KIR, and CD57 define a process of CD56 dim NK-cell differentiation uncoupled from NK-cell education. Blood 2010; 116: 3853–3864.

37 Nielsen CM, White MJ, Goodier MR, Riley EM. Functional Significance of CD57 Expression on Human NK Cells and Relevance to Disease. Front Immunol 2013; 4: 422.

38 Boyiadzis M, Memon S, Carson J, Allen K, Szczepanski MJ, Vance BA et al. Up-regulation of NK Cell Activating Receptors Following Allogeneic Hematopoietic Stem Cell Transplantation under a Lymphodepleting Reduced Intensity Regimen is Associated with Elevated IL-15 Levels. Biol Blood Marrow Transplant 2008; 14: 290–300.

39 Deaglio S, Zubiaur M, Gregorini A, Bottarel F, Ausiello CM, Dianzani U et al. Human CD38 and CD16 are functionally dependent and physically associated in natural killer cells. Blood 2002; 99: 2490–2498.

40 Zambello R, Barilà G, Manni S, Piazza F, Semenzato G. NK cells and CD38: Implication for (Immuno)Therapy in Plasma Cell Dyscrasias. Cells 2020; 9: 768.

41 Erokhina SA, Streltsova MA, Kanevskiy LM, Telford WG, Sapozhnikov AM, Kovalenko EI. HLA-DR+ NK cells are mostly characterized by less mature phenotype and high functional activity. Immunol Cell Biol 2018; 96: 212–228.

42 Yamabe T, Takakura K, Sugie K, Kitaoka Y, Takeda S, Okubo Y et al. Induction of the 2B9 antigen/dipeptidyl peptidase IV/CD26 on human natural killer cells by IL-2, IL-12 or IL-15. Immunology 1997; 91: 151–158.

43 McKinney EF, Cuthbertson I, Harris KM, Smilek DE, Connor C, Manferrari G et al. A CD8+ NK cell transcriptomic signature associated with clinical outcome in relapsing remitting multiple sclerosis. Nat Commun 2021; 12: 635.

44 Solomon KR, Chan M, Finberg RW. Expression of GPI-Anchored Complement Regulatory Proteins CD55 and CD59 Differentiates Two Subpopulations of Human CD56+CD3-Lymphocytes (NK Cells). Cell Immunol 1995; 165: 294–301.

45 Siebert N, Jensen C, Troschke-Meurer S, Zumpe M, Jüttner M, Ehlert K et al. Neuroblastoma patients with high-affinity FCGR2A, -3A and stimulatory KIR 2DS2 treated by long-term infusion of anti-GD 2 antibody ch14.18/CHO show higher ADCC levels and improved event-free survival. Oncoimmunology 2016; 5: e1235108.

46 Cheung NK V., Cheung IY, Kushner BH, Ostrovnaya I, Chamberlain E, Kramer K et al. Murine anti-GD2 monoclonal antibody 3F8 combined with granulocyte-macrophage colony-stimulating factor and 13-cis-retinoic acid in high-risk patients with stage 4 neuroblastoma in first remission. J Clin Oncol 2012; 30: 3264–3270.

47 Delgado DC, Hank JA, Kolesar J, Lorentzen D, Gan J, Seo S et al. Genotypes of NK cell KIR receptors, their ligands, and Fcγ receptors in the response of neuroblastoma patients to Hu14.18-IL2 immunotherapy. Cancer Res 2010; 70: 9554–9561.

48 Tarek N, Luduec JB Le, Gallagher MM, Zheng J, Venstrom JM, Chamberlain E et al. Unlicensed NK cells target neuroblastoma following anti-GD2 antibody treatment. J Clin Invest 2012; 122: 3260–3270.

49 Giebel S, Dziaczkowska J, Czerw T, Wojnar J, Krawczyk-Kulis M, Nowak I et al. Sequential recovery of NK cell receptor repertoire after allogeneic hematopoietic SCT. Bone Marrow Transplant 2010; 45: 1022–1030.

50 Béziat V, Descours B, Parizot C, Debré P, Vieillard V. NK Cell Terminal Differentiation: Correlated Stepwise Decrease of NKG2A and Acquisition of KIRs. PLoS One 2010; 5: e11966.

51 Brenchley JM, Karandikar NJ, Betts MR, Ambrozak DR, Hill BJ, Crotty LE et al. Expression of CD57 defines replicative senescence and antigen-induced apoptotic death of CD8+ T cells. Blood 2003; 101: 2711–2720.

52 Shifrin N, Raulet DH, Ardolino M. NK cell self tolerance, responsiveness and missing self recognition. Semin Immunol 2014; 26: 138–144.

53 Thielens A, Vivier E, Romagné F. NK cell MHC class I specific receptors (KIR): From biology to clinical intervention. Curr Opin Immunol 2012; 24: 239–245.

54 Elliott JM, Yokoyama WM. Unifying concepts of MHC-dependent natural killer cell education. Trends Immunol 2011; 32: 364–372.

55 Vento-Tormo R, Efremova M, Botting RA, Turco MY, Vento-Tormo M, Meyer KB et al. Single-cell reconstruction of the early maternal–fetal interface in humans. Nature 2018; 563: 347–353.

56 Koopman LA, Kopcow HD, Rybalov B, Boyson JE, Orange JS, Schatz F et al. Human Decidual Natural Killer Cells Are a Unique NK Cell Subset with Immunomodulatory Potential. J Exp Med 2003; 198: 1201–1212.

57 Farley MJ, Bartlett DB, Skinner TL, Schaumberg MIAA, Jenkins DG. Immunomodulatory Function of Interleukin-15 and Its Role in Exercise, Immunotherapy, and Cancer Outcomes. Med Sci Sports Exerc 2023; 55: 558–568.

58 Huntington ND, Legrand N, Alves NL, Jaron B, Weijer K, Plet A et al. IL-15 trans-presentation promotes human NK cell development and differentiation in vivo. J Exp Med 2009; 206: 25–34.

59 Carson WE, Giri JG, Lindemann MJ, Linett ML, Ahdieh M, Paxton R et al. Interleukin (IL) 15 is a novel cytokine that activates human natural killer cells via components of the IL-2 receptor. J Exp Med 1994; 180: 1395–1403.

60 Castriconi R, Dondero A, Bellora F, Moretta L, Castellano A, Locatelli F et al. Neuroblastoma-Derived TGF-β1 Modulates the Chemokine Receptor Repertoire of Human Resting NK Cells. J Immunol 2013; 190: 5321–5328.

61 Zenarruzabeitia O, Vitallé J, Astigarraga I, Borrego F. Natural killer cells to the attack: Combination therapy against neuroblastoma. Clin Cancer Res 2017; 23: 615–617.

62 Tie Y, Tang F, Peng D, Zhang Y, Shi H. TGF-beta signal transduction: biology, function and therapy for diseases. Mol Biomed 2022; 3: 45.

63 Bottino C, Dondero A, Bellora F, Moretta L, Locatelli F, Pistoia V et al. Natural Killer Cells and Neuroblastoma: Tumor Recognition, Escape Mechanisms, and Possible Novel Immunotherapeutic Approaches. Front Immunol 2014; 5: 56.

64 Kleinertz H, Hepner-Schefczyk M, Ehnert S, Claus M, Halbgebauer R, Boller L et al. Circulating growth/differentiation factor 15 is associated with human CD56bright natural killer cell dysfunction and nosocomial infection in severe systemic inflammation. EBioMedicine 2019; 43: 380–391.

65 Albonici L, Giganti MG, Modesti A, Manzari V, Bei R. Multifaceted Role of the Placental Growth Factor (PlGF) in the Antitumor Immune Response and Cancer Progression. Int J Mol Sci 2019; 20: 2970.

66 Autiero M, Luttun A, Tjwa M, Carmeliet P. Placental growth factor and its receptor, vascular endothelial growth factor receptor-1: Novel targets for stimulation of ischemic tissue revascularization and inhibition of angiogenic and inflammatory disorders. J Thromb Haemost 2003; 1: 1356–1370.

67 Fredriksson L, Li H, Eriksson U. The PDGF family: Four gene products form five dimeric isoforms. Cytokine Growth Factor Rev 2004; 15: 197–204.

68 Andrae J, Gallini R, Betsholtz C. Role of platelet-derived growth factors in physiology and medicine. Genes Dev 2008; 22: 1276–1312.

69 Lokker NA, Sullivan CM, Hollenbach SJ, Israel MA, Giese NA. Platelet-derived growth factor (PDGF) autocrine signaling regulates survival and mitogenic pathways in glioblastoma cells: Evidence that the novel PDGF-C and PDGF-D ligands may play a role in the development of brain tumors. Cancer Res 2002; 62: 3729–3735.

70 Servidei T, Riccardi A, Sanguinetti M, Dominici C, Riccardi R. Increased sensitivity to the platelet-derived growth factor (PDGF) receptor inhibitor STI571 in chemoresistant glioma cells is associated with enhanced PDGF-BB-mediated signaling and STI571-induced Akt inactivation. J Cell Physiol 2006; 208: 220– 228.

71 Ma S, Tang T, Wu X, Mansour AG, Lu T, Zhang J et al. PDGF-D−PDGFRβ signaling enhances IL-15–mediated human natural killer cell survival. Proc Natl Acad Sci 2022; 119: e2114134119.

72 Keskin DB, Allan DSJ, Rybalov B, Andzelm MM, Stern JNH, Kopcow HD et al. TGFβ promotes conversion of CD16+ peripheral blood NK cells into CD16-NK cells with similarities to decidual NK cells. Proc Natl Acad Sci U S A 2007; 104: 3378–3383.

73 Cerdeira AS, Rajakumar A, Royle CM, Lo A, Husain Z, Thadhani RI et al. Conversion of Peripheral Blood NK Cells to a Decidual NK-like Phenotype by a Cocktail of Defined Factors. J Immunol 2013; 190: 3939–3948.

74 Hawke LG, Mitchell BZ, Ormiston ML. TGF-β and IL-15 Synergize through MAPK Pathways to Drive the Conversion of Human NK Cells to an Innate Lymphoid Cell 1–like Phenotype. J Immunol 2020; 204: 3171–3181.

75 Du X, Zhu H, Jiao D, Nian Z, Zhang J, Zhou Y et al. Human-Induced CD49a+ NK Cells Promote Fetal Growth. Front Immunol 2022; 13: 821542.

76 Crinier A, Milpied P, Escalière B, Piperoglou C, Galluso J, Balsamo A et al. High-Dimensional Single-Cell Analysis Identifies Organ-Specific Signatures and Conserved NK Cell Subsets in Humans and Mice. Immunity 2018; 49: 971–986.e5.

77 Cruz-Zárate D, Cabrera-Rivera GL, Ruiz-Sánchez BP, Serafín-López J, Chacón-Salinas R, López-Macías C et al. Innate Lymphoid Cells Have Decreased HLA-DR Expression but Retain Their Responsiveness to TLR Ligands during Sepsis. J Immunol 2018; 201: 3401–3410.

78 Liem LM, Fibbe WE, van Houwelingen HC, Goulmy E. Serum transforming growth factor-beta1 levels in bone marrow transplant recipients correlate with blood cell counts and chronic graft-versus-host disease. Transplantation 1999; 67: 59–65.

79 Siewiera J, Gouilly J, Hocine H-R, Cartron G, Levy C, Al-Daccak R et al. Natural cytotoxicity receptor splice variants orchestrate the distinct functions of human natural killer cell subtypes. Nat Commun 2015; 6: 10183.

80 Kielsen K, Oostenbrink LVE, von Asmuth EGJ, Jansen-Hoogendijk AM, van Ostaijen-ten Dam MM, Ifversen M et al. IL-7 and IL-15 Levels Reflect the Degree of T Cell Depletion during Lymphopenia and Are Associated with an Expansion of Effector Memory T Cells after Pediatric Hematopoietic Stem Cell Transplantation. J Immunol 2021; 206: 2828–2838.

81 Wischhusen J, Melero I, Fridman WH. Growth/Differentiation Factor-15 (GDF-15): From Biomarker to Novel Targetable Immune Checkpoint. Front Immunol 2020; 11: 951.

82 Gonzalez VD, Huang Y-W, Delgado-Gonzalez A, Chen S-Y, Donoso K, Sachs K et al. High-grade serous ovarian tumor cells modulate NK cell function to create an immune-tolerant microenvironment. Cell Rep 2021; 36: 109632.

83 Albini A, Noonan DM. Decidual-Like NK Cell Polarization: From Cancer Killing to Cancer Nurturing. Cancer Discov 2021; 11: 28–33.

84 Gallazzi M, Baci D, Mortara L, Bosi A, Buono G, Naselli A et al. Prostate Cancer Peripheral Blood NK Cells Show Enhanced CD9, CD49a, CXCR4, CXCL8, MMP-9 Production and Secrete Monocyte-Recruiting and Polarizing Factors. Front Immunol 2021; 11: 586126.

85 Bruno A, Bassani B, D’Urso DG, Pitaku I, Cassinotti E, Pelosi G et al. Angiogenin and the MMP9-TIMP2 axis are up-regulated in proangiogenic, decidual NK-like cells from patients with colorectal cancer. FASEB J 2018; 32: 5365–5377.

86 Heldin CH, Westermark B. Mechanism of action and in vivo role of platelet-derived growth factor. Physiol Rev 1999; 79: 1283–1316.

87 Erokhina SA, Streltsova MA, Kanevskiy LM, Grechikhina M V., Sapozhnikov AM, Kovalenko EI. HLA-DR-expressing NK cells: Effective killers suspected for antigen presentation. J Leukoc Biol 2021; 109: 327–337.

88 Wang L, Halliday D, Johnson PM, Christmas SE. Expression of complement regulatory proteins on human natural killer cell subsets. Immunol Lett 2007; 112: 104–109.

89 Liu Y, Beyer A, Aebersold R. On the Dependency of Cellular Protein Levels on mRNA Abundance. Cell 2016; 165: 535–550.

90 Cenik C, Cenik ES, Byeon GW, Grubert F, Candille SI, Spacek D et al. Integrative analysis of RNA, translation, and protein levels reveals distinct regulatory variation across humans. Genome Res 2015; 25: 1610–1621.

91 Funaro A, Reiniš M, Trubiani O, Santi S, Di Primio R, Malavasi F. CD38 Functions Are Regulated Through an Internalization Step. J Immunol 1998; 160: 2238–2247.

92 Lam JKP, Azzi T, Hui KF, Wong AMG, McHugh D, Caduff N et al. Co-infection of Cytomegalovirus and Epstein-Barr Virus Diminishes the Frequency of CD56dimNKG2A+KIR− NK Cells and Contributes to Suboptimal Control of EBV in Immunosuppressed Children With Post-transplant Lymphoproliferative Disorder. Front Immunol 2020; 11: 1–12.

93 Lopez-Sejas N, Campos C, Hassouneh F, Sanchez-Correa B, Tarazona R, Pera A et al. Effect of CMV and Aging on the Differential Expression of CD300a, CD161, T-bet, and Eomes on NK Cell Subsets. Front Immunol 2016; 7: 476.

94 Campos C, Pera A, Sanchez-Correa B, Alonso C, Lopez-Fernandez I, Morgado S et al. Effect of age and CMV on NK cell subpopulations. Exp Gerontol 2014; 54: 130–137.

95 Pradier A, Simonetta F, Waldvogel S, Bosshard C, Tiercy J-M, Roosnek E. Modulation of T-bet and Eomes during Maturation of Peripheral Blood NK Cells Does Not Depend on Licensing/Educating KIR. Front Immunol 2016; 7: 299.

96 Muntasell A, Servitja S, Cabo M, Bermejo B, Pérez-Buira S, Rojo F et al. High numbers of circulating CD57+ NK cells associate with resistance to her2-specific therapeutic antibodies in HER2+ primary breast cancer. Cancer Immunol Res 2019; 7: 1280–1292.

97 Jungraithmayr W, Enz N. CD26 – The emerging role of a costimulatory molecule in allograft rejection. Cell Mol Immunol 2020; 17: 1208–1209.

98 Drakul M, Čolić M. Immunomodulatory activity of dipeptidyl peptidase-4 inhibitors in immune-related diseases. Eur J Immunol 2023; 53: e2250302.

99 Morimoto C, Schlossman SF. The structure and function of CD26 in the T-cell immune response. Immunol Rev 1998; 161: 55–70.

100 Kopinska A, Krawczyk-Kulis M, Dziaczkowska-Suszek J, Bieszczad K, Jagoda K, Kyrcz-Krzemien S. The importance of the number of transplanted cells with dipeptidyl peptidase-4 expression on the haematopoietic recovery and lymphocyte reconstitution in patients with multiple myeloma after autologous haematopoietic stem-cell transplantation. Hematol Oncol 2017; 35: 225–231.

101 Kandekar S, Punatar S, Khattry N, Gokarn A, Jindal N, Mirgh S et al. Low levels of CD26 on certain cellular subtypes of donor harvest is associated with better clinical outcomes post allogeneic stem cell transplantation through regulation of NF-κB pathway and pro-inflammatory cytokines. Int Immunopharmacol 2023; 125: 111054.

102 Finberg RW, White W, Nicholson-Weller A. Decay-accelerating factor expression on either effector or target cells inhibits cytotoxicity by human natural killer cells. J Immunol 1992; 149: 2055–2060.

103 Han M, Hu L, Wu D, Zhang Y, Li P, Zhao X et al. IL-21R-STAT3 signalling initiates a differentiation program in uterine tissue-resident NK cells to support pregnancy. Nat Commun 2023; 14: 7109.

